# Coordinated function of paired NLRs confers *Yr84*-mediated stripe rust resistance in wheat

**DOI:** 10.1101/2024.06.14.599041

**Authors:** Valentyna Klymiuk, Krystalee Wiebe, Harmeet Singh Chawla, Jennifer Ens, Rajagopal Subramaniam, Curtis J. Pozniak

## Abstract

Cloning of resistance genes expands our understanding of their function and facilitates their deployment in breeding. Here, we report the cloning of two genes from wild emmer wheat (*Triticum turgidum* ssp. *dicoccoides*) underlying *Yr84*-mediated stripe rust resistance using a combination of fine mapping, long read-sequencing and mutation-induced functional validation. In contrast to all previously cloned stripe rust genes, the incompletely dominant *Yr84* phenotype is conferred through the coordinated function of paired nucleotide-binding leucine-rich repeats (NLR) genes *CNL* and *NL*. We hypothesize that based on their genomic organization, annotation, expression profiles and predicted protein structure, CNL functions as a sensor NLR (sNLR) responsible for effector recognition, and NL acts as a helper NLR (hNLR) initiating downstream resistance cascades. The CNL and NL lack an integrated domain(s) previously implicated in effector recognition by paired NLRs, therefore these findings contribute new insights into plant paired NLRs structure and molecular mechanisms of function.

## Introduction

Wheat is a cornerstone of global agriculture and a vital staple in human diets worldwide, providing 20% of the calories consumed by humans. (Shiferaw et al. 2013). Bread wheat (*Triticum aestivum*; 2n=6x=42; AABBDD) and durum wheat (*T. turgidum* ssp. *durum*; 2n=4x=28, AABB) dominate global production, while less-known subspecies like spelt (*T. spelta*; 2n=6x=AABBDD), emmer wheat (*T. turgidum* subsp. *dicoccum*; 2n=4x=48; AABB), Khorasan or Kamut (*T. turgidum* ssp. *turanicum*; 2n=4x=28; AABB) and einkorn (*T. monococcum*; 2n=2x=14; AA) occupy niche growth regions, but still hold significant agricultural value.

Wheat production faces challenges from biotic stresses, including various pathogens and pests, which thrive under favorable climate conditions and spread through wind, machinery, and human activity. Among these, wheat stripe (yellow) rust, caused by the fungus *Puccinia striiformis* f. sp. *tritici* (*Pst*), is one of the most significant biotic challenges. Its global prevalence poses a recurring risk to wheat production, with outbreaks showing both regional variability and cyclical patterns. Under epidemic conditions, yield losses can reach as high as 100% for susceptible varieties. The dynamic evolution of this pathogen underscores the ongoing challenge of combating this disease. For example, the emergence of new races from sexually recombining populations in the center of stripe rust diversity (Hovmøller et al. 2016) illustrates this ongoing threat.

Genetic resistance remains the most effective strategy for mitigating yield losses caused by wheat stripe rust, prompting extensive research to identify resistance genes in the primary, secondary, and tertiary gene pools of wheat. The catalogue of mapped stripe rust genes comprises more than 130 formally and temporary designated *Yr* genes (McIntosh et al. 2020), but only 11 have been cloned so far: *Yr36* (Fu et al. 2009), *Yr18*(*Lr34*) (Krattinger et al. 2009), *Yr10* (Liu et al. 2014), *Yr46*(*Lr67*) (Moore et al. 2015), *Yr15* (Klymiuk et al. 2018), *Yr5*/*YrSp* (Marchal et al. 2018), *Yr7* (Marchal et al. 2018), *YrAS2388* (Zhang et al. 2019), *YrU1* (Wang et al. 2020), *Yr27* (Athiyannan et al. 2022), *YrNAM* (Ni et al. 2023). However, wheat-pathogen co-evolution often leads to the eventual breakdown of widely deployed resistance genes. To mitigate this, best practice involves pyramiding multiple *Yr* genes into a single genetic background, achievable through marker-assisted selection (MAS) using tightly-linked or gene-specific markers. Thus, discovery, functional validation and deployment of additional resistance is needed to expand the repertoire of available genes for effective stripe rust management.

The majority of cloned disease resistance genes fall into two main classes of R proteins: nucleotide-binding leucine-rich-repeat (NLR) proteins (Baggs et al. 2017) and Tandem Kinase Proteins (TKP) (Klymiuk et al. 2021). Among these, NLRs represent a larger and more extensively studied class. They are members of the Signal-Transduction ATPases with Numerous Domain (STAND) superfamily, and function as signaling proteins to initiate immune responses in infected tissues. The architecture of NLR genes is notably diverse, typically characterized by a central NB-ARC domain (Nucleotide Binding site and ARC) coupled to multiple Leucine Rich Repeats (LRR). At the N-terminus, they may contain a coiled-coil (CC; CNL subfamily), Resistance to Powdery Mildew 8 (RPW8; RNL subfamily) or toll/interleukin 1 receptor (TIR; TNL subfamily) domains (Chia and Carella 2023). Additionally, they may carry noncanonical integrated domains (ID) of various structures, including heavy-metal-associated domain (HMA), WRKY, NO3-induced (NOI)/RIN4, BED-type zinc finger domain, kinases, or others, which can appear as N- or C-terminal domains (Grund et al. 2019). Studies have shown that these IDs are involved directly or indirectly in effector recognition (Cesari et al. 2014) and may serve as effector decoys (Xi et al. 2022). While in most cases, singleton NLRs are generally sufficient for pathogen recognition and immune response, functional studies have confirmed that some NLRs act in pairs – one as a sensor (sNLR) which recognizes the pathogen, and the second as a helper (hNLR) which executes resistance cascades (Xi et al. 2022).

Wild emmer wheat (*T. turgidum* ssp. *dicoccoides* 2n=4x=28, AABB), a wild progenitor of cultivated bread and durum wheats, grows across a diverse range of natural habitats in the Fertile Crescent and has accumulated a wide range of adaptations to various growth conditions (Nevo et al. 2002). This natural variation positions wild emmer wheat as a valuable source of novel alleles lost through domestication and plant breeding, particularly for disease resistance traits (Huang et al. 2016). Furthermore, the relative ease of crossing wild emmer wheat with both durum and bread wheat enhances the utility of these novel alleles in breeding. A genome assembly of wild emmer wheat accession Zavitan is available to support genetic and genomic studies (Avni et al. 2017) but gene cloning remains challenging due to suppressed recombination, presence/absence variation (PAV) and structural variations (Li et al. 2024). However, the declining cost of long-read sequencing technologies has enabled rapid assembly and annotation of complex genomes, characterization of genome structures and variations (Athiyannan et al. 2022) making gene cloning increasingly feasible. Concurrently, RNA sequencing technologies, such as IsoSeq facilitate annotation and detection of alternative isoforms and accurate delineation of transcription start sites, polyadenylation sites, and alternative splicing events (Zhang et al. 2022).

Previously, we identified an accession of wild emmer wheat highly resistant to wheat stripe rust and confirmed that resistance was conferred by a single, incompletely dominant gene (*Yr84*) that we localized using BSA-Seq (Klymiuk et al., 2022). In the current study, we further applied a long-read whole genome sequencing and mutation-based functional validation strategy to demonstrate the combined function of paired NLR genes *CNL* and *NL* in conferring *Yr84*-mediated stripe rust resistance. Based on detailed analysis of gene expression patterns, protein conserved motifs, and protein-protein interaction study, our data supports that CNL likely functions as an sNLR, while NL is responsible for initiating resistance cascades. Haplotype analysis in a diverse collection of tetraploid wheat *T. turgidum* species indicated the uniqueness of these functionally co-dependent *CNL* and *NL* genes in wild emmer wheat, from which we successfully transferred resistance into durum and bread wheat cultivars using the validated *CNL* gene-specific KASP marker.

## Results

### Fine mapping and *Yr84* candidate genes

Previously, we used a BSA-Seq approach to localize *Yr84* to a 2.2 cM interval on chromosome 1BS of tetraploid wheat (Klymiuk et al. 2022). To more precisely localize *Yr84*, we screened 1039 F_2_ plants with *Yr84*-flanking markers *usw310* and *usw318* and identified 56 putative recombinants. Subsequently, these were phenotyped and genotyped with additional KASP markers (*usw310, usw312, usw313, usw314, usw315, usw316, usw317* and *usw318*) to precisely identify positions of recombination events (Table S1) and further delineate the *Yr84* region. From these 56, 20 F_2_ plants representing distinct recombination events were advanced to develop F_2:3_ homozygous recombinants. These were again phenotyped and genotyped with *usw310, usw312*-*usw317* markers. This reduced the *Yr84* interval to 0.5 cM, flanked by *usw313* and *usw316* markers and co-segregated with *usw314* and *usw315* (Figure 1A, Table S2). The physical region spanning *usw313* and *usw316* corresponds to 975 Kb (chr1B: 10967359-11942601) in the Zavitan v2 assembly (Figure 1B). This reduced the previously published list of *Yr84* candidate genes in the Zavitan assembly (Klymiuk et al. 2022) from 79 to 22 (Table S3), of which 11 were annotated as high confidence genes encoding proteins involved in disease resistance (Table S3, in bold).

**Figure 1.**
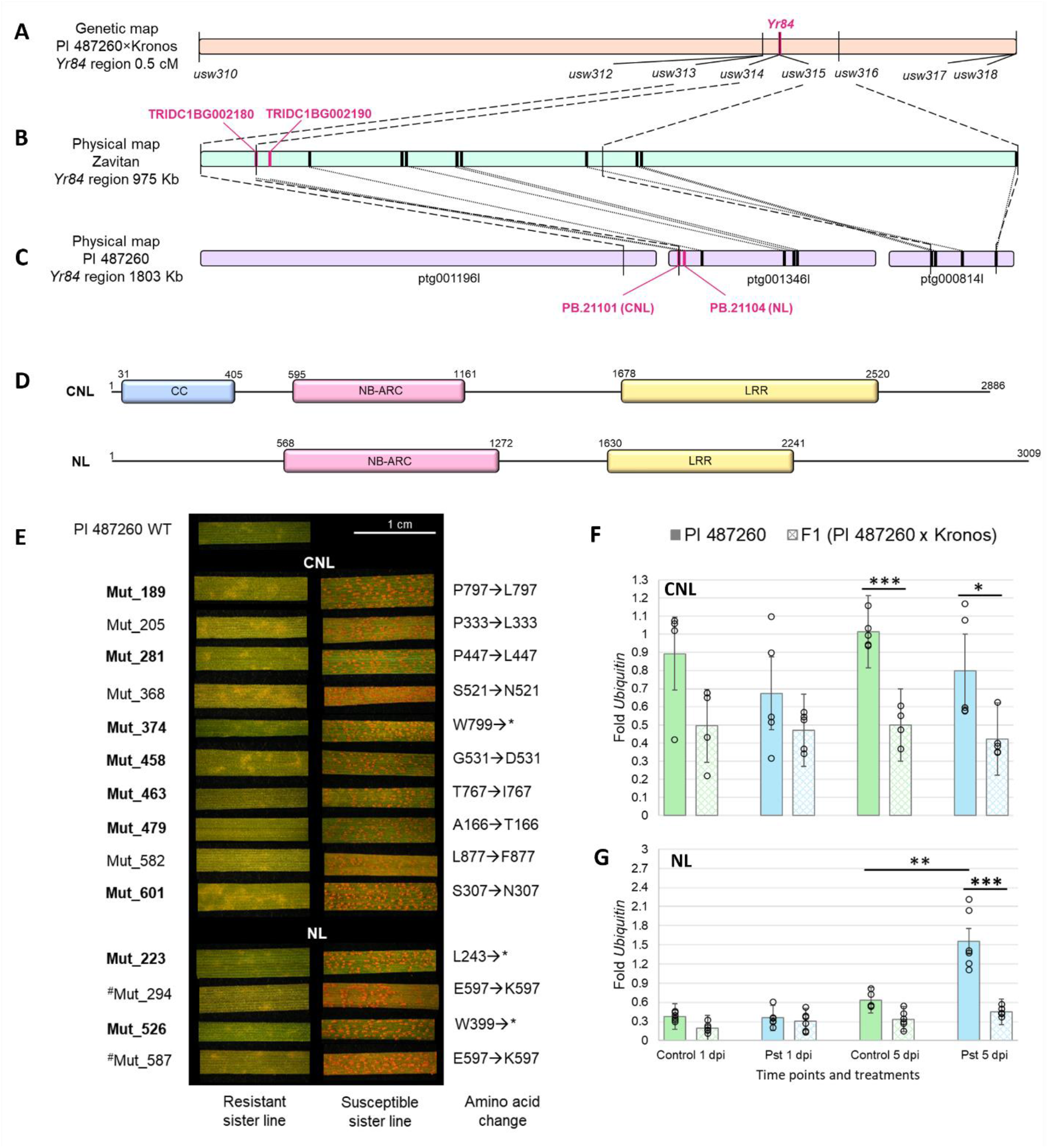
Fine mapping of *Yr84* and functional validation of candidate genes. (**A-C**) Fine mapping of the *Yr84* gene: (**A**) genetic map based on screening of 1039 F_2_ plants from the cross Kronos x PI 487260. (**B**) *Yr84* physical interval in Zavitan v1 assembly, 11 genes annotated to be involved in disease resistance responses are indicated as black rectangles, (**C**) *Yr84* physical interval in PI 487260 assembly represented by three contigs, positions of the genes homologous to those identified in the Zavitan are collinear. (**D**) *CNL* and *NL* genomic structure and domain architecture. (**E**) Phenotypic changes in mutants carrying mutations in *CNL* or *NL* genes: homozygous mutants exhibit a susceptible response to *Pst* inoculation, while their sister lines from the same mutant families carrying no mutations show a resistance response. Amino acid changes of predicted protein sequence are noted, with * indicating a premature stop codon; (**F-G**) *CNL* (**F**) and *NL* (**G**) expression studies: CNL expression is stable in both, PI 487260 and F_1_ plants (PI 487260 x Kronos); *NL* expression is elevated at 5 dpi in PI 487260, but not in F_1_ plants. Bars are SD, while asterisks denote significant differences between the treatments according to a one-tailed *t*-test: *P*<0.05 (*), *P*<0.01 (**), P<0.001 (***).

### Identification and validation of *Yr84* candidate genes

To fully characterize the gene space and to facilitate mutational analysis of *Yr84*, we first developed a low coverage, long read assembly of the wild emmer wheat accession PI 48726 using 11X whole genome generated PacBio HiFi CCS-reads. The final assembly comprised 10,46 Mbp assembled into 4,615 contigs. The N_50_ of the assembly was 6.7 Mbp, with an L_50_ of 440 (Table S4). Comparative analysis with the published short-read assembly of Zavitan (Avni et al. 2017) (Table S4) revealed that the PI 487260 assembly was 465 Mbp longer (Table S4). BUSCO metrics indicated the presence of 99% complete BUSCO genes (4853 out of 4896 BUSCO genes), supporting the that the CCS assembly sufficiently captured the gene content (Figure S1). We anchored the *Yr84* interval to the PI 487260 assembly (Figure 1C) and identified four contigs spanning 1.8 Mbp. This was nearly twice the size observed in Zavitan, so we compared it against available PacBio long-read assemblies of hexaploid wheat Fielder (Sato et al. 2021) and Kariega (Athiyannan et al. 2022) and determined the *Yr84* region in these assemblies spans 979 Kbp and 960 Kbp, respectively. In contrast, the interval in tetraploid wheat cv. Svevo (V.2), spanned 1.4 Mbp. Comparison to available hexaploid wheat assemblies from the 10+ Genome Project (Walkowiak et al. 2020) revealed the *Yr84* region spanned from 950 Kbp to 1.4 Mbp. These discrepancies suggest that *Yr84* likely resides in a hyper-variable region of chromosome 1B. Despite these discrepancies, the number and order of genes in the *Yr84* interval were largely conserved between Zavitan and the PI 487260 low coverage assembly (Figure 1B,C, Table S3).

To identify genes underlying the *Yr84* phenotype, we screened 754 M_2_ EMS-derived mutant families of PI 487260 over the course of two independent screening experiments. In the first experiment, nine susceptible mutants were identified for which we generated high-quality whole genome Illumina paired-end short reads (Table S5). The number of paired reads generated ranged from 330,022,753 to 791,863,544 representing 5-13X coverage per genotype (Table S5). We mapped the Illumina reads to the PI 487260 PacBio assembly to identify mutations in the delineated *Yr84* region. This analysis revealed that seven susceptible mutants harbored independent mutations in a CC-NB-ARC-LRR (*CNL*) gene (Figure 1D,E), while the remaining two exhibited mutations in a NB-ARC-LRR (*NL*) gene (Figure 1D,E). KASP markers were developed based on the detected mutations (Table S6) and these confirmed the presence of homozygous mutations in susceptible mutants and the presence of the resistant allele in corresponding resistant sister lines and WT PI 487260 (Figure S2).

In a second screening experiment, we identified an additional five susceptible mutants. For these we performed ONT-based amplicon sequencing of the *CNL* and *NL* genes. This resulted in >50X read coverage per mutant. Three susceptible mutants were identified to harbor mutations in the *CNL* gene when compared to the PI 487260 assembly, while the remaining two exhibited mutations in the *NL* gene. Thus, based on both screens, we identified a total of fourteen independent mutations – ten in the *CNL* gene and four in the *NL* gene (Figure 1E). These results confirm that these genes are functionally co-dependent, and both are required for *Yr84*-mediated stripe rust resistance. Additional EMS-derived mutations in CNL and NL together with functional complementation and knock out assays in an independent genetic background confirm our results (Hu et al. 2024). This represents the first reported case of paired-NLRs conferring resistance to stripe rust in wheat.

### *CNL* and *NL* expression, structure and conserved motifs

Having confirmed that both *CNL* and *NL* genes were responsible for the *Yr84*-mediated resistance phenotype, we examined their relative expression and performed ISO-Seq based RNA sequencing to support their annotation (Table S7). Repeated experiments confirmed that the *CNL* gene was constitutively expressed, while *NL* was upregulated in the presence of the pathogen (*P*<0.01) (Figure 1F-G). Additionally, we observed that the *CNL* and *NL* expression levels in heterozygous F_1_ plants at 5 dpi were generally lower and did not follow the same trend as the homozygous carrier line PI 487260 (*P*<0.001) (Figure 1G). Isoform detection revealed 17 isoforms for the *CNL* gene (PB.21101.1-PB.21101.17) and 7 isoforms for the *NL* gene (PB.21104.1-PB.21104.7) (Figures S3-S4). For the *CNL* gene, all isoforms except PB.21101.1, PB.21101.3, PB.21101.15, PB.21101.16 are predicted to encode identical proteins (Figure S3). The translation of the remaining four isoforms results in truncated proteins. For the *NL* gene, all isoforms except PB.21104.6 and PB.21104.7 encode nearly identical protein sequences, with the exception of isoforms PB.21104.2 and PB.21104.4, which have an extra amino acid at position 4 of the protein sequence, not affecting the domain architecture (Figure S4). The remaining two isoforms are disrupted and likely result in non-functional proteins.

Given these data, we annotated the genomic sequence of the *CNL* gene to span 3511 bp (ptg001346l:44183-47693), resulting in a 2886 bp coding sequence. Based on an NCBI conserved domain search, the *CNL* gene comprises CC (31-405 positions of the CDS sequence), NB-ARC (595-1161 bp), and LRR (1678-2520) domains (Figure 1D). The genomic sequence of *NL* spans 3310 bp (ptg001346l:74784-78093), corresponding to 3009 bp coding sequence. The *NL* includes a predicted NB-ARC domain (568-1272 positions of the CDS sequence) and LRR (1630-2241) (Figure 1D). No IDs or transmembrane domains were detected in either the *CNL* or *NL* genes.

Next, we performed a detailed assessment of conserved motifs and subdomains within the predicted CNL and NL protein sequences to assess conservation and predict functionality (see Supplementary Text 1 for details). This analysis revealed the presence of NBD^VG^, NBD^P-loop^, NBD^Walker-B^, NBD^RNSB-B^, ARC1^RNSB-C^, ARC1^GLPL^, ARC2^RNSB-D^, ARC2^MHD^ and multiple LRRLxxLxL motifs for both proteins (Figure 2A, Figures S5, S6). Additionally, the CNL protein exhibited conserved CC^EDVID^ and NBD^RNSB-A^ motifs which were absent in the NL (Figure 2A, Figure S5). We detected NBD subdomain motifs VG (involved in nucleotide contact), P-loop (responsible for phosphate-binding), RNBS-A (essential for nucleotide binding site formation), Walker-B (required for Mg2+ ion coordination for ATPase activity) and RNBS-B (ligand binding), and these were found to be highly conserved in the CNL protein but only moderately conserved in the NL protein (Figure 2A, Supplementary Text 1). Likewise, amino acid conservation in MHD (nucleotide-binding and regulation of subdomain interactions), located in the ARC2 subdomain, exhibited extensive variability in both CNL and NL (Figure 2A, Supplementary Text 1) and is likely only functional in the CNL protein. An analysis of the presence of the arginine residues from the LRR^R-cluster^ motif showed that both CNL and NL are conserved only in the first three aa positions (marked with red triangles on Figure 2A), while the remaining two are variable (marked with yellow triangles on Figure 2A). Analysis of the positions of our EMS-induced mutations relative to protein structure indicated that all were loss-of-function mutations, with some affecting the NBL^RNBS-D^ and LRR^LxxLxL^ conserved motifs (Figure 2A). However, the majority of mutations were localized outside of the conserved motifs, likely affecting general protein 3D structure.

**Figure 2.**
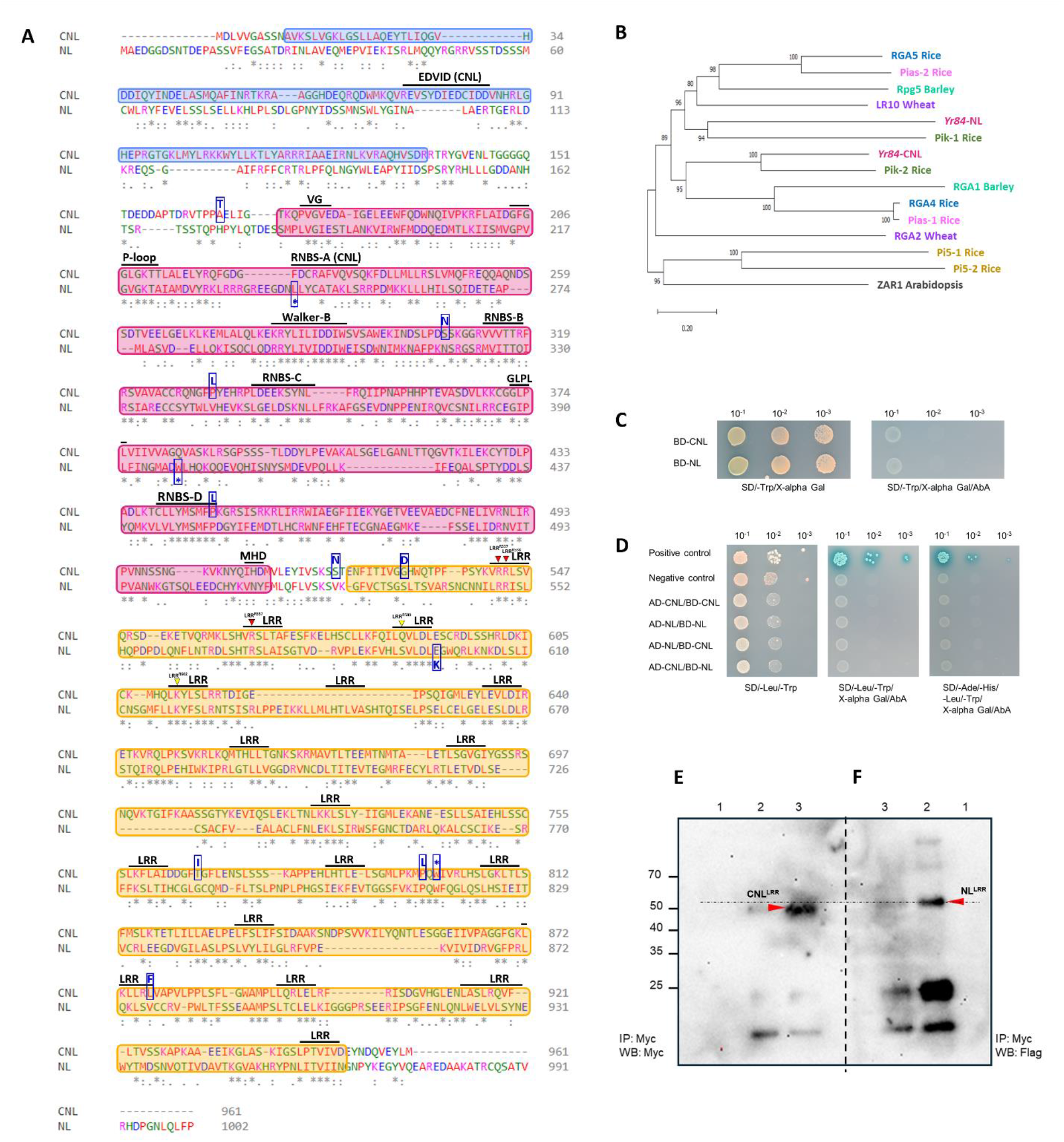
CNL and NL as paired NLRs. (**A**) CNL and NL protein alignment. Colored boxes denote approximate borders for CC (blue), NB-ARC (pink), and LRR (yellow) domains defined based on conserved motifs. Black lines indicate positions of conserved motifs, with their names labeled. Red triangles mark conserved Arginine from LRR^R-cluster^, where applicable in CNL/NL proteins, while yellow triangles indicate where this conservation is disrupted. Position labels are based on the SR35 LRR^R-cluster^ (Förderer et al. 2022). Blue boxes denote positions of identified loss-of-function mutations, with corresponding amino acid change or stop codon (*) specified within the box. (**B**) Phylogenetic tree of CNL/NL and other paired NLRs discovered in cereals. NLRs from the same pairs are presented in the same colors: RGA5/RGA4 (blue), Pias-1/Pias-2 (pink), Rpg5/RGA1 (green), LR10/RGA2 (violet), Pik-1/Pik-2 (olive), Pi5-1/Pi5-2 (gold), CNL/NL (mulberry). ZAR1 was used as an outgroup. (**C-D**) Yeast two-hybrid screening showing no interactions between full-length CNL and NL proteins. (**C**) Autoactivation test demonstrating no autoactivation of either bait proteins; (**D**) Mating test indicating no interaction for CNL/CNL, NL/NL, and CNL/NL pairs. (**E-F**) Co-IP experiments showing interaction between CNL^LRR^ and NL^LRR^. *N. benthamiana* leaves were co-infiltrated with empty vectors (PGW511 and PGW517; *Lane 1*) and co-infiltrated with CNL^LRR^-Myc tagged and NL^LRR^-Flag tagged genes. Proteins were immunoprecipitated with anti-Myc-magnetic microbeads for 30 mins (*Lane 2*) and overnight (*Lane 3*): (**E**) Western blot with anti-Myc antibodies showed a positive signal for CNL^LRR^ with ∼ molecular weight of 49 kD. (**F**) The Western blot with anti-Flag antibodies show interaction of CNL^LRR^ with NL^LRR^ (∼ molecular weight 52 kD). The red arrows indicate the migration of each protein respective to each other (horizontal dashed lines). The vertical dashed lines indicate the two independent Western blots from the same the gel. IP: Immunoprecipitation; WB: Western blot.

Given the presence of two co-dependent NLR genes underlying the *Yr84* phenotype, we hypothesized they may act as a functional, interacting pair. We examined the protein similarity of known cereal paired NLRs and determined that CNL/NL were most similar to rice Pik-1/Pik-2 (Figure 2B) – a well-studied model system for plant paired NLRs (Ashikawa et al. 2008; Zhai et al. 2014; Zdrzałek et al., 2020). Like Pik-1/Pik-2 and other paired-NLRs, CNL and NL are physically linked (<27 Kbp) and arranged in a head-to head orientation (Figure 1C). Using the analogy of physically interacting paired NLRs (Zhai et al. 2014; Huh et al. 2017), we assessed the potential interaction between CNL and NL at the protein level using Y2H and coimmunoprecipitation (Co-IP) assays. The Y2H screen revealed no interaction between full-length CNL/NL, NL/NL and CNL/CNL proteins when these two proteins are co-expressed in yeast cells (Figure 2C,D). Both CNL and NL were tested as baits, but the results were consistent – no interaction in yeast for CNL/NL, NL/NL and CNL/CNL protein pairs. However, it has been shown that LRR domains can interact in a ligand-independent manner to form, for example, LRR-based cell surface interaction network (CSI^LRR^) (Smakowska-Luzan et al. 2018). Therefore, we conducted Co-IP experiments to test for interaction between CNL^LRR^ and NL^LRR^, and found that they indeed interact *in vivo* (Figure 2E,F). Co-IP with anti-Myc antibodies, followed by probing with anti-Myc antibodies, showed the correct size of CNL^LRR^ (∼49 kD). The same Co-IP experiment, immunoblotted with anti-Flag antibodies, revealed a protein of ∼52kD, the expected size for NL^LRR^, suggesting an interaction between the two proteins.

### *CNL* and *NL* diversity

We conducted a search for *CNL* and *NL* copies in multiple wheat genome assemblies, including Zavitan (Zhu et al. 2019), Svevo (Maccaferri et al. 2019), Chinese Spring (IWGSC 2018), and CDC Landmark (Walkowiak et al. 2020). The search revealed *CNL* copies on chromosomes 1B, 2B, and 5D of these reference genomes (Figure S7). The *CNL* copies were found in three contigs of CCS assembly of PI 487260, corresponding to chromosomes 1B, 2B and 5B (Figure S7). In contrast, the *NL* copy was only identified on 1B chromosomes of all screened assemblies, and a single contig of the low coverage PI 487260 assembly (Figure S7).

Previously we recommended the KASP marker *usw314* as a tool for marker assisted selection of *Yr84* (Klymiuk et al. 2022). Based on comparative analysis of available wheat genome assemblies to the PI 487260 assembly reported here, this marker discriminates between resistant PI 487260 allele and multiple susceptible alleles. Thus, we utilized this marker to characterize two tetraploid wheat diversity panels (Klymiuk et al. 2023; Mazzucotelli et al. 2020). This revealed that the PI 487260 *CNL* allele was absent in all cultivated and domesticated tetraploid wheat accessions. Only 38 genotypes (21%) of wild emmer wheat showed the presence of the PI 487260 *CNL* allele, of which 31 exhibited susceptible responses to inoculation with *Pst*. Thus, we comprehensively assess the sequence variation in these 38 accessions by amplicon sequencing, which revealed SNPs and INDELs in the exons of both *CNL* and *NL* genes. For some accessions we did not obtain amplification of one of the genes (marked as N/A in Table S8), likely due to substantial sequence differences at the primer position(s) for amplification (Table S6). In total, we identified ten protein coding haplotypes of the *CNL* gene, and fifteen haplotypes of the *NL* gene based on differences within exons (Tables S8-10). We did not observe a clear trend in the prevalence of haplotype combinations based on geographic distribution, except that all genotypes collected in Lebanon shared the same haplotypes for both genes (*CNL* Hap2 and *NL* Hap12). The co-occurrence of specific CNL and NL haplotypes appears to be independent, suggesting an ancient diversification event (Table S8).

Six of the 38 accessions showed a resistance response to *Pst* inoculation; however, five of them carried the highly effective *Yr15* (*WTK1*) gene based on functional marker genotyping (Klymiuk et al. 2019). However, one accession, PI 471699, lacked *Yr15,* and exhibited resistance to *Pst.* Additionally, it carried unique haplotypes for both *CNL* (Hap10; Table S9) and *NL* (Hap15 Table S10) genes, suggesting that these could in fact represent alternative resistant alleles of *Yr84*. To test this hypothesis, we crossed PI 487260 with PI 471699 and developed an F_2_ population. A resistant phenotype was observed for the F_1_ plant, and the F_2_ population exhibited a phenotypic segregation ration of 15:1 for resistant (R) to susceptible (S) individuals in response to *Pst* inoculation (χ^2^ = 0.33; *P* = 0.05), suggesting duplicate gene action from the presence of two independently assorting resistance genes. Additionally, we developed KASP markers to differentiate between the *CNL* and *NL* alleles in these two WEW accessions: *usw329* marker in *CNL* gene and *usw330* marker in the *NL* gene (Table S6). The analysis revealed that variation in *CNL*/*NL* did not associate with resistance (Table S11) and we conclude that PI 471699 carries non-functional alleles of *CNL*/*NL*, and that the resistance in that line is conferred by resistance gene(s) other than *Yr84*.

### *Yr84* introgression into cultivars

While both *CNL*/*NL* genes are essential for *Yr84*-mediated resistance, their close physical proximity (∼27 Kbp) would allow for the simultaneous introgression into breeding material using a single KASP marker. We applied the *CNL* gene-specific KASP marker *usw314* to introgress *Yr84* into several Canadian durum and bread wheat cultivars (Figure 3). In all cases, *Yr84* conferred improved resistance. Upon introgression, previously susceptible bread wheat cv CDC Landmark, BW1085, as well as moderately resistant durum wheat cv CDC Precision and cv CDC Evident, achieved full resistance against stripe rust (Figure 3). Also, as resource to the wheat community, we introgressed *Yr84* into the Australian cultivar Avocet, a universally susceptible background cultivar for the stripe rust differential set used globally for pathogenicity tests. Introgression of the *CNL*/*NL* pair into Avocet conferred the expected *Yr84* resistance phenotype (Figure 3).

**Figure 3.**
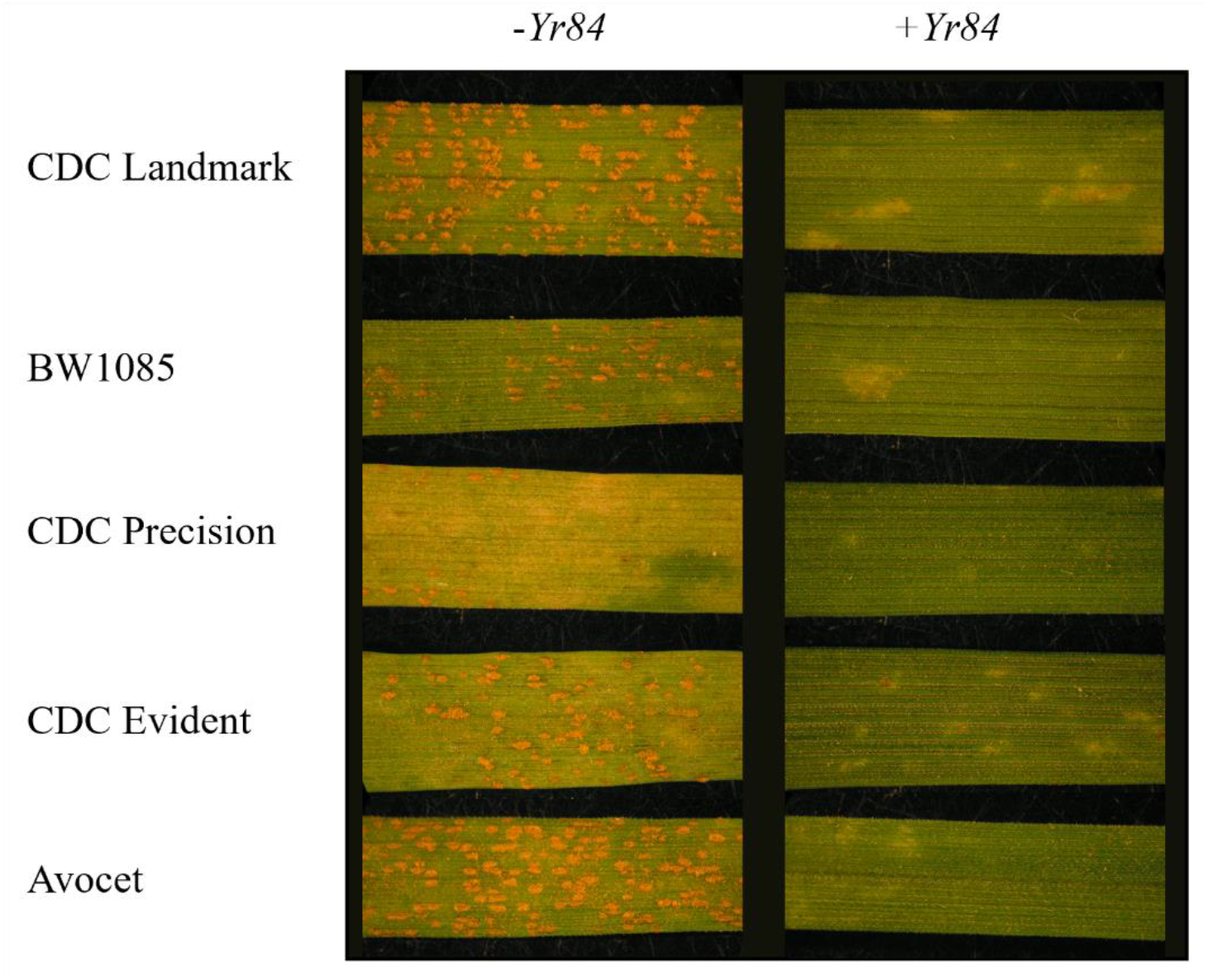
Phenotypic responses of original cultivars and breeding lines and same lines carrying *Yr84* introgression. Phenotyping performed with *Pst* W001 race. CDC Landmark, BW1085 and Avocet are *T. aestivum* lines, all susceptible to W001. CDC Precision and CDC Evident are *T. turgidum* ssp. *durum* lines, both moderately resistant to W001.

## Discussion

Discovery and cloning of resistance genes remain crucial for ensuring sustainable wheat production by facilitating the incorporation of novel sources of resistance into elite varieties to combat emerging pathogen strains. Over the past decade, rapid advancements in cloning methods and genome sequencing technologies have revolutionized resistance gene cloning (R. Chen et al. 2024). The advent of long-read sequencing technologies, such as PacBio and ONT, has significantly reduced the cost, making it more practical to sequence large, complex genomes, like that of wheat (Yu et al. 2023). However, the cost of generating reference-quality pseudomolecule assemblies still remains considerable and is not necessary to support gene cloning. In this study, we utilized low-coverage (11X coverage) PacBio whole-genome sequencing to obtain a contig-level assembly of the *Yr84*-carrier wild emmer wheat accession PI 487160. This approach was sufficient to assemble the gene space, that together with our high-resolution mapping data enabled rapid cloning of the genes underlining the *Yr84* phenotype. The combination of this assembly along the IsoSeq transcriptomic data generated in this study adds to the genomic resources already available to support gene cloning in wheat.

NLR genes represent the most abundant and well-known class of resistance genes and many have been functionally validated to provide resistance against a wide range of plant pathogens and pests. While most cloned NLRs function independently, some NLRs function as pairs to confer resistance. To date, a number of paired NLRs have been functionally validated across diverse plant species: *Arabidopsis* (RRS1/RPS4, RRS1B/RPS4B, RPP2A/RPP2B, CHS1/SOC3, CSA1/CHS3), melon (Fom-1/Prv), rapeseed *Brassica napus* (BnRPR1/BnRPR2), rice (RGA4/RGA5, Pik-1/Pik-2, Pi5-1/Pi5-2 or Pii-1/Pii-2, Pias1/Pias2), barley (RGA1/Rpg5), and wheat (LR10/RGA2, CNL1/CNL5) (reviewed by Xi et al. 2022; Zhang Y et al. 2017; Yang et al. 2022; Mermigka et al. 2023; Zhu et al. 2024). Paired NLRs are common, and genome studies across various plant species have revealed that ∼20% of NLRs may function as pairs (Chen et al. 2021; L. Wang et al. 2019). In our and companion work (Hu et al. 2024), EMS mutagenesis, gene silencing, CRISPR/Cas9-mediated genome editing knock-out and complementation via transformation all confirmed that the *CNL* and *NL* are both required for *Yr84*-mediated stripe rust resistance. This work represents the first reported cased on paired-NLRs conferring resistance to the wheat stripe rust pathogen.

The *Yr84 CNL*/*NL* pair share some common features of typical paired-NLRs. They are physically linked and are oriented in a head-to-head confirmation (Xi et al. 2022). Within one-to-one functional pairs, one partner is typically involved in pathogen effector recognition (sensor NLR, sNLR), while the other signals downstream resistance cascades (helper NLR, hNLR) (Xi et al. 2022). Typical sNLRs contain an integrated domain which is proposed to form the bases for the molecular mechanism of function, serving as effector-targeted decoys (Xi et al. 2022). These integrated domains may include WRKY, RATX, NOI, TRX, HMA, NCKX among others (J. Wang et al. 2019; Zhu et al. 2024). However, effector binding may not be limited to the integrated domain, but could include a combination of multiple interfaces, including a C-terminal Lys-rich stretch following the integrated domain, as demonstrated for the rice RGA5 protein (X. Zhang et al. 2024). Interestingly (and novel to our work), neither the CNL protein nor the NL protein of the *Yr84* pair possess integrated domains, while both carry an LRR domain known to be important for the specificity of effector recognition (Ma et al. 2020; Martin et al. 2020). Our expression studies suggest that the *CNL* likely serves as sNLR in the *Yr84* NLR pair. We observed that the *CNL* is constitutively expressed, and its expression remains unchanged in the presence of the pathogen. Conversely, *NL* gene expression was upregulated at 5 dpi, supporting its role as a hNLR. These results are consistent with expression of *Pikh-1* (sNLR) which is constitutively expressed, while *Pikh-2* (hNLR) expression increases ≈2-fold 3 dpi (Zhai et al. 2014. Interestingly, *NL* gene expression in heterozygous F_1_ plants from PI 487160xKronos cross was not upregulated in response to pathogen attack and generally both, *CNL* and *NL* expression levels were lower in F_1_ plants compared with PI 487260. We hypothesize that this discrepancy may contribute to the incomplete dominance of *Yr84* (Klymiuk et al. 2022) where the lower expression and possibly abundance of NL protein in these plants might not be sufficient to confer full resistance.

The sequence of conserved motifs within paired NLR proteins can provide insights into their on-to-one functionality. For example, in the CC-NB-ARC protein of the stem rust resistance gene Sr35, arginines of the LRR^R-cluster^ form salt-bridges with acidic residues in the CC^EDVID^ motif, which are crucial for proper intramolecular packing (Förderer et al. 2022). Additionally, the EDVID motif’s direct interaction with the LRR domain is essential for both autoinhibition and activation of the ZAR1 protein (Burdett et al. 2019). Given the conservation of the LRR^R-cluster^ in both the CNL and NL proteins, a similar interaction between the EDVID motif of the CNL protein and LRR domain(s) of CNL or NL may be pivotal for their function. While we did detect a CC domain-unique EDVID motif in the NL protein sequence, its level of conservation was low (Figure 2A, Supplementary Text 1), leading us to hypothesize that NL lacks a functional CC domain. Given that our understanding of the structure and function of diverse CC domains is still evolving (Bentham et al. 2018), this question may need to be revisited in the future. We also have observed that the ARC2^MHD^ motif of the NL protein diverges from conserved sequences (Figure 2A, Supplementary Text 1): Histidine (H), the most conserved amino acid within this motif (Van Ooijen et al. 2008), is replaced by Asparagine (N) in the NL sequence (Figure 2). The MHD motif plays a crucial regulatory role in the activity of R proteins, with involvement in coordinating nucleotide binding and controlling subdomain interactions (Van Ooijen et al. 2008) and studies have shown that such amino acid replacement results in autoactivation (Van Ooijen et al. 2008). Thus, it is reasonable to hypothesize that the NL protein alone might exhibit autoactivity and it is suppressed by its CNL pair. Concurrently, the interaction of CNL with the pathogen effector is likely to alter its confirmation, leading to the release of NL activity, resulting in hypersensitive response. However, Hu et al. (2024) demonstrated that HR was not observed in Tobacco leaves overexpressing NL alone in the absence of the pathogen effector. This suggests that the CNL/NL pair may function similarly to Pikp-1/Pikp-2 through helper-sensor cooperation rather than through negative regulation (Zdrzałek et al. 2020). On the other hand, our Y2H and Co-IP experiments revealed that although no interaction of the full-length proteins was detected in Y2H experiments, their LRR domains showed interaction when co-expressed in Tobacco leaves. Moreover, it has been shown that CC domains of the CNL proteins self-associate (Hu et al. 2024) suggesting formation of homo-complex previously described for Pikp-1/Pikp-2 paired NLRs (Zdrzałek et al. 2020). Thus, while the precise molecular function of CNL/NL pair remains to be uncovered, ligand-independent interaction at the LRR domain level within this pair has been confirmed.

From an evolutionary perspective, both CNL and NL genes exhibited multiple susceptible alleles in wild emmer wheat natural populations (Tables S8-10). Despite extensive screening of over 170 wild emmer wheat accessions, we were unable to detect a haplotype identical to the resistance-haplotype discovered in PI 487260. Hu et al. (2024) also did not detect the *Yr84* haplotypes while screening 910 *Triticum* accessions but did identify other rare resistant haplotypes for both genes: *NLR1*, which differs from the *CNL* reported here at two amino acid changes (S520L and A866T), and *NLR2* which corresponds to *NL*, but carries one amino acid change (E110K) (Hu et al. (2024). This uniqueness of resistant R gene alleles has been described for the wild emmer wheat powdery mildew resistance genes *Pm69* (Li et al. 2024) and *MlIW170*/*Pm26* (Zhu et al., 2024), but contrasts with the situation for the wild emmer wheat stripe rust resistance gene *Yr15*, whose resistant allele was found in 16% of populations (Klymiuk et al. 2019).

We have successfully developed *Yr84* introgression lines, which have been integrated into Canadian durum and bread wheat breeding programs. Furthermore, we have developed the Avocet+*Yr84* line and propose its inclusion into the stripe rust differential set. This addition will facilitate comprehensive assessments of the durability of the *Yr84* against diverse stripe rust isolates from various geographic regions. Given incomplete dominance of the *Yr84*, its introgression requires the use of Marker-Assisted-Selection (MAS). To this end, we recommend employing the *CNL* gene-specific KASP marker *usw314* (Klymiuk et al. 2022). The strategic integration of *Yr84* into elite wheat cultivars underscores its potential to improve stripe rust resistance and contribute to sustainable wheat production globally.

## Materials and Methods

### Plant material

We previously localized *Yr84* from a tetraploid wild emmer wheat accession PI 487260 (SY 20121), which was obtained from the USDA-ARS National Small Grains Collection through GRIN-Global (https://npgsweb.ars-grin.gov/gringlobal/search). Kronos (https://www.goldenstategrains.com/crop-variety/kronos), which is susceptible to the stripe rust race used in this study, was used to develop an F_2_ population with PI 487260. The seed of M_1_ EMS-treated PI 487260 plants was provided by Prof. Tzion Fahima, University of Haifa, Israel. CDC Landmark (spring hexaploid wheat), CDC Precision (durum wheat; (Pozniak & Clarke 2016)) and CDC Evident (durum wheat) are cultivars, while BW1085 (spring hexaploid wheat) is a breeding line, all developed at the Crop Development Centre, University of Saskatchewan, Canada. *Yr84* was backcrossed four times into these lines using *Yr84*-carrier lines KSY_10-64 (Kronos/PI 487260) and RSY_2-2-1-7 (Ruta/PI 487260//3*Ruta) (Klymiuk et al. 2022) as the source of *Yr84*: Landmark+*Yr84* (CDC Landmark/RSY-2-2-1-7//3*CDC Landmark), BW1085+*Yr84* (BW1085/RSY-2-2-1-7//3*BW1085), CDC Precision+*Yr84* (CDC Precision/KSY-10-64//3*CDC Precision), CDC Evident+*Yr84* (CDC Evident/KSY-10-64//3*CDC Evident). We also introgressed *Yr84* into the Australian bread wheat cv Avocet (Avocet+*Yr84*) by crossing Avocet with PI 487260, followed by five additional backcrosses to Avocet (Avocet/PI 487260//6*Avocet). The seed of a tetraploid wheat diversity panel, comprising wild emmer wheat (177 accessions), domesticated emmer wheat (131 genotypes), and durum wheat landraces (139 genotypes) was obtained from previous study (Klymiuk et al. 2023). The seed of subset of 95 genotypes (Table S12) from the Global Durum Wheat Panel (GDP) (Mazzucotelli et al. 2020) were received from ICARDA.

### Phenotyping and genotyping

Populations were phenotyped for resistance to stripe rust using the W001 *Pst* race provided by Dr. Randy Kutcher from the University of Saskatchewan, as detailed in (Klymiuk et al. 2022). The W001 is virulent on *YrA*, *Yr2*, *Yr6*, *Yr7*, *Yr8*, *Yr9*, *Yr17*, *Yr25*, *Yr27*, *Yr28*, *Yr29*, *Yr31*, *YrSu* (Brar et al. 2018). In brief, seedlings were inoculated at the two-leaf stage were with *Pst* uredinia spores mixed with mineral oil, left to air dry at room temperature for 1h, and then placed in a humidity chamber set at 99% relative humidity for 24h. Subsequently, the plants were moved to a growth chamber set at 23C with 16h of light, 8 hours of dark. Stripe rust infection types (scale 0-9; 0-3 being resistant, 4-6 moderately resistant and 7-9 susceptible; Line & Qayoum, 1992) were recorded at 14 days post inoculation (dpi).

The *usw310*-*usw318* markers were developed in a previous study, and all KASP genotyping assays were conducted as described in (Klymiuk et al. 2022). Primers for amplicon sequencing, and expression studies of *CNL* and *NL* candidate genes were developed (Table S6) from annotated gene sequences obtained from a PacBio circular consensus sequence (CCS) assembly of PI 487260. All primers underwent BLAST analysis against the PI 487260 and Zavitan (WEWSeq version 2.0 assembly; Zhu et al. 2019) genome assemblies to confirm their chromosome specificity.

### PI 487260 PacBio assembly

To clone the genes responsible for *Yr84*-derived resistance, we developed a low-coverage, PacBio CCS assembly of PI 487260. High molecular weight (HMW) DNA was extracted from approximately 1.5 g young leaf tissue (genotype: PI 487260) using the Nucleobond HMW DNA extraction protocol (Macherey Nagel 740160.20). The HMW DNAs were quantified by fluorometry (Qubit 2.0), and DNA integrity was assessed using a Tapestation 2200 instrument (Agilent, Santa Clara, California, US). Subsequently, the HMW DNAs were stored at 4°C until library preparation. A total of 25µg of HMW DNA was sent to KeyGene (NV, Wageningen, the Netherlands) for library preparation and sequencing. HiFi data were generated from a total of six SMRTcells on a Sequel IIe instrument (Pacific Biosystems). This resulted in approximately 126Gb of HiFi data, achieving 11X genome coverage with an average read length of approximately 16kb (mean number of passes per molecule = 12). Raw CCS (consensus circular sequencing) reads were obtained in BAM format and converted to fastq using the bam2fastq tool (version 1.3.0) from PacBio. A *de novo* assembly was generated using hifiasm v.0.15.4-r347 with default parameters. Assembly statistics were estimated using QUAST version 5.0.2. Assessment of genomic data quality was performed using BUSCO (Manni et al. 2021) and by assessment of synteny of the *Yr84* genomic region between PI 487260 and the published Zavitan WEWSeq version 2.0 assembly (Zhu et al. 2019).

### Fine mapping and candidate genes

The F_2_ plants from the Kronos x PI 487260 mapping population were screened with flanking markers *usw310* and *usw318* (Klymiuk et al. 2022) to identify recombinants. The recombinant F_2_ plants were then screened with previously co segregating markers (*usw310, usw312-usw317*) to further delineate the region and identify independent recombination events, which were advanced to the next generation. These F_3_ families were phenotyped with *Pst* and screened with *usw310* and *usw312*-*usw317* markers to select homozygous recombinants. Physical positions of newly identified flanking markers *usw313* and *usw316* were located in Zavitan WEWSeq version 2.0 assembly (Zhu et al. 2019) to identify the narrowed *Yr84* interval. High confidence genes resining within this region were extracted based on gene models of Zavitan genome assembly. The *Yr84* physical region was also identified in Svevo v1 (Maccaferri et al. 2019), Svevo v2 (https://wheat.pw.usda.gov/GG3/content/released-triticum-turgidum-durum-wheat-svevo-rel-20-pseudomolecules), cv Fielder (v1; Sato et al. 2021), cv Kariega (v1; Athiyannan et al. 2022) and assemblies of LongReach Lancer, ArinaLrFor, Jagger, SY Mattis and Julius (Walkowiak et al. 2020) using *usw313* and *usw316* flanking markers.

### PI 487260 Iso-Seq

Total RNA was isolated from the center of the second leaf of *Pst*-non-inoculated PI 487260 seedling at two-leave stage using the RNeasy plant mini kit (Qiagen). A non-size-selected PacBio Iso-Seq library was prepared and sequenced on a Sequel II system (PacBio) with a 30-hour movie length (v10) on one SMRTcell by Genome Quebec (Montreal, Canada). The raw BAM file with CCS reads was processed to obtain the isoforms and transcript structures using the SMRTLink version 10 pipeline from PacBio. In brief, the reads were demultiplexed and the sequencing primers were removed using the LIMA tool (version 2.1.0). Chimeric and truncated reads were then removed using the IsoSeq3 refine tool. The full-length non-chimeric (FLNC) reads were used to call isoforms using IsoSeq3 cluster. The isoforms were subsequently aligned to the long-read assembly of PI 487260 using pbmm2 and finally collapsed to generate high-quality isoforms using IsoSeq3 collapse.

### Development of susceptible *Yr84* mutants

The M_0_ seed was obtained by mutagenizing PI 487260 using 0.3% ethyl methanesulfonate (EMS) solution following published protocol (Klymiuk et al. 2018). The M_1_ plants were grown under greenhouse conditions for seed increase. In total, 12-20 seeds from each M_2_ family were sown into 5×10 cell trays (with two cells per each family and 6-10 seeds per cell). Non-mutagenized PI 487260 control and susceptible control Avocet S were included with two replicates in each tray. The trays were watered and placed in a constant 10°C cold chamber/fridge for 3-5 days to promote uniform germination, after which they were transferred to growth chamber with a temperature of 23/18°C and light regime of 16/8 h.

Plants at two-leave stage (approximately 9-10 days old) were inoculated following the procedure summarized above and described in (Klymiuk et al. 2022). Screening of the plants was conducted at 14 dpi and M_2_ families showing segregation for resistance to stripe rust race W001 were selected as candidate mutant families. One of the most susceptible (susceptible sister line) and one of the most resistant (resistant sister line) plants from each candidate family were transplanted and advanced to the M_3_ generation. DNA was extracted from each of the selected M_2_ plants for subsequent sequencing and genotyping.

In total, 20 M_3_ embryos from each resistant and susceptible sister line from each candidate mutant family were subjected to the same phenotypic screening procedure to confirm their phenotypic response. Binocular pictures of phenotypic responses were recorded for all sister lines from all mutant families. A summary of phenotypic responses of confirmed susceptible mutants is presented in Figure 1E).

### PI 487260mut sequencing and detection of mutations in *CNL* and *NL* genes

Individual paired-end libraries were prepared from the M_2_ generation for each susceptible sister line from nine independent M_2_ families (Mut_189, Mut_223, Mut_281, Mut_374, Mut_458, Mut_463, Mut_479, Mut_526, Mut_601) using the Illumina DNA Prep protocol (Illumina PN 20018704). Libraries were uniquely indexed for multiplexing using IDT for Illumina Nextera DNA UD and quantified by qPCR (Kapa Biosystems). The libraries were normalized, pooled into two pools consisting of 4 and 5 libraries per pool, and the final DNA concertation and library size was estimated using a Tapestation 2200 (Agilent). Each pool was sequenced across one lane of NovaSeq 6000 S4 PE150 to target approximately 10X coverage per genotype. Sequencing data were demultiplexed and aligned to the PI 487260 PacBio CCS reference assembly using bwa.

Additionally, five mutants (Mut_205, Mut_368, Mut_582, Mut_294, Mut_587) were amplicon sequenced using Oxford Nanopore Technologies (ONT) using procedures we developed previously (Nilsen et al. 2020). The amplicons were obtained by amplifying PI 487260 DNA with primers located 37-138 bp outside of *CNL*/*NL* gene sequence using LongAmp® Hot Start Taq 2X Master Mix (New England Biolabs, Ipswich, Massachusetts, US) following the manufacturer’s protocol. The presence of PCR product was confirmed on a 1% agarose gel, and positive samples were used for ONT library preparation using Native Barcoding Kit 96 V14, following manufacturer’s protocol (Oxford Nanopore Technologies, Oxford, United Kingdom). The pooled library was sequenced using one flow-cell on PromethION (Oxford Nanopore Technologies). The obtained reads were aligned to *CNL* and *NL* PI 487260 genomic sequences using minimap2, followed by visualization of alignments using Integrative Genomic Viewer (IGV).

### *CNL* and *NL* architecture and copies in reference genomes

NCBI Conserved Domain Search (https://www.ncbi.nlm.nih.gov/Structure/cdd/wrpsb.cgi) and InterProScan tool (https://www.ebi.ac.uk/interpro/search/sequence/) were utilized to identify the domain architecture of the *CNL* and *NL* genes in the PI 487260 genome. Protein sequences were predicted from Iso-Seq sequences using the Expasy Translate tool (https://web.expasy.org/translate/). Conserved motifs within each of the CNL and NL proteins were detected using NLRexpress (https://nlrexpress.biochim.ro; (Martin et al. 2022). Positions of LRR^R-cluster^ (Förderer et al. 2022) were identified by alignment of CNL and NL sequences with SR35 protein (Förderer et al. 2022). An alignment of CNL and NL proteins was conducted using Clustal Omega at EMBL-EBI (https://www.ebi.ac.uk/Tools/msa/clustalo/). BLASTn was employed to search for *CNL* and *NL* copes in reference genomes Zavitan (WEWSeq version 2.0 assembly; (Zhu et al. 2019), Svevo (Durum Wheat Svevo Rel. 1.0 pseudomolecules; Maccaferri et al. 2019), Chinese Spring (IWGSC RefSeq v1.0; IWGSC 2018) and CDC Landmark (PGSBv2.1; Walkowiak et al. 2020). *CNL* copies from PI 487260 contig assembly were assigned to specific chromosomes based on their best blast hits in Chinese Spring pseudomolecule reference genome. The phylogenetic tree of all *CNL* and *NL* copies was visualized using EMBL-EBI Simple Phylogeny online tool (https://www.ebi.ac.uk/Tools/phylogeny/simple_phylogeny/).

### Expression studies

Sampling of plant tissues for expression studies was conducted during the phenotypic testing using race W001. Both PI 487260 and F_1_ plants (PI 487260×Kronos) were subjected to inoculation with *Pst* (mineral oil + *Pst* spores) or mock (control) inoculation (mineral oil only), as detailed in (Klymiuk et al. 2022). Two time points were chosen for each treatment: 1 dpi and 5 dpi, which were selected based on preliminary analysis on a smaller experimental set. Sampling at each time point for each treatment was performed from six biological replicates for *NL* gene and five biological replicates for *CNL* gene. Tissues were sampled from the middle of the second leaf, and total RNA was extracted using the RNeasy Plant Mini Kit (Qiagen).

To assess gene expression, we developed gene-specific *NL* and *CNL* primers to amplify short fragments of each gene (Table S6). The primers were blasted against the PI 487260 assembly and Zavitan assembly to ensure their specificity to the allele of interest. Preliminary tests demonstrated no amplification with these primers from Kronos RNA, confirming their usability to detect expression of the functional allele in F_1_ plants (PI 487260 x Kronos).

Quantitative PCR (qPCR) was performed in a 96-well plate using a CFX96 Touch Real-Time PCR Detection System (Bio-Rad Laboratories, Hercules, CA, US) with Bio-Rad EvaGreen qPCR Supermix assay. Three technical replicates were performed for each biological replicate, and *Ubiquitin* was used as a reference gene. An analysis of variance and one-tailed *t*-tests were applied to assess the statistical significance (*P*<0.05) of differences in expression patterns detected within this experiment.

### Paired NLR phylogeny

The protein sequences of functionally validated paired-NLRs from rice (RGA4/RGA5 (Okuyama et al. 2011), Pik-1/Pik-2 (Ashikawa et al. 2008), Pi5-1/Pi5-2 (Lee et al. 2009), Pias1/Pias2 (Shimizu et al. 2022), barley (RGA1/Rpg5 (Wang X et al. 2013), and wheat (LR10/RGA2 (Lourte et al. 2009) were collected from NCBI. The NLR protein HOPZ-ACTIVATED RESISTANCE 1 (ZAR1; AT3G50950) from Arabidopsis was used as an outgroup. Full-length proteins were aligned for phylogenetic analysis using MEGA Software (www.megasoftware.net/) and the phylogenetic tree was constructed using Neighbor-joining method with 10,000 bootstraps and visualized employing MEGA Software.

### Yeast two-hybrid (Y2H) assay

We conducted a Y2H assay to assess the potential interaction of *Yr84* associated CNL/NL proteins. Specifically, we used a Matchmaker Gold Yeast Two-Hybrid System (Takara Bio Inc., cat.

\# 630489) to explore interactions between NL/NL, CNL/CNL and NL/CNL pairs, following the manufacturer’s protocol. The coding sequences (CDS) corresponding to predicted CNL and NL protein sequences were synthesized by General Biosystems (Durham, NC, United States) and subsequently used for PCR amplification with PrimeSTAR® Max DNA Polymerase assay (Takara Bio Inc., cat. # R045A) for cloning into pGBKT7 DNA-BD and pGADT7 AD vectors (Table S6). Yeast transformation was carried out using the Yeastmaker™ Yeast Transformation System 2 (Takara Bio Inc., cat. # 630439).

### Co-Immunoprecipitation (Co-IP) assays

The protein sequences of CNL^LRR^ and NL^LRR^ used for Co-IP assays are provided in Supplementary Text 2. CNL^LRR^ was cloned into the gateway expression vector pGW517 with Myc tag at the C-terminus. NL^LRR^ was cloned into pGW511 with Flag tag at the C-terminus. The clones were transformed into the *Agrobacterium* strain LBA4404. The two genes were transiently expressed in *N. benthamiana* plants and leaves were harvested after 50 hrs and flash frozen in liquid N2 (Ma et al., 2012). Proteins were extracted with lysis buffer (50 mM Tris·HCl (pH 7.5), 50 mM NaCl, 1% Triton X-100, 5 mM DTT and protease inhibitor cocktail (Sigma). Following incubation on ice for 30 mins, coimmunoprecipitation experiments were performed with 750 µg of protein extract and incubated with anti-Myc (cat# 13-2500, Thermofisher, Canada) magnetic microbeads (Miltenyi Biotec) were used to pull down the proteins associated with CN^LRR^ tagged Myc protein complex. The beads were washed two times with 50 mM Tris·HCl (pH 7.5), 50 mM NaCl, 1% Triton X-100 and once with 50 mM Tris·HCl (pH 7.5), 50 mM NaCl and an equal aliquot was analyzed by SDS-PAGE and Western blot analysis. Both Myc-tagged (CNL^LRR^) and Flag tagged (NL^LRR^) proteins were detected with 1/5,000 dilution of the anti-Myc and anti-Flag (cat# MA1-91878, Thermofisher) primary antibodies, followed by HRP-conjugated goat-anti mouse secondary antibodies at 1/20,000 dilution (Goat anti-mouse-HRP conjugate, cat# 31430, Thermofisher). The proteins were detected by SuperSignal chemiluminescent substrate (Thermo Fisher). The images were processed in the iBright CL750 imaging system (Invitrogen).

### Diversity panel screening and sequencing

The *usw314* marker, previously developed in *Yr84* mapping region (Klymiuk et al. 2022), was found to reside within the *CNL* gene. It was subsequently employed to screen a tetraploid wheat diversity panel (Klymiuk et al. 2023). A subset of wild emmer wheat accessions from this panel were selected for ONT amplicon-based re-sequencing of the two genes to identify variation in sequence and structure of their *Yr84*-associated *CNL* and *NL* alleles. Out of 39 selected accessions, 36 carried a putative functional allele of the *CNL* gene, two were heterozygous, and Zavitan was selected as *Yr84* non-carrier control. The amplicon sequencing procedure was identical to that described above for the five PI 487260 mutants.

## Data availability

The PacBio CCS reads (SRR28097670-SRR28097686) and IsoSeq data (SRR28097687) for PI 487260 accession have been deposited to NCBI under BioProject PRJNA1079375. The PI 487260 contig assembly and Iso-Seq transcript structures have been deposited to Zenodo (Klymiuk et al., 2024). Functional PI 487260 *CNL* and *NL* sequences have been deposited to NCBI under accession numbers PP841906 and PP841907, respectively.

## Acknowledgement

The authors are thankful to Prof. Tzion Fahima, University of Haifa, for providing M_1_ seed of PI 487260 used in the current study, which allowed us to screen for segregating families to select and characterize susceptible mutants. The authors are grateful to Thomas Kroj (INRAE) and Matthew Moscou (USDA-ARS) for fruitful scientific discussions and to Madison Kist, Ayla Lichtenwald, Xue (Snow) Lin, Lexie Gerl, and Maureen Troesch for their technical support.

## Author contributions

Jointly supervised research: V.K., C.J.P.; conceived and designed the experiments: V.K., K.W., H.S.C., J.E., R.S., C.J.P.; performed the experiments: V.K., K.W., H.S.C., J.E., R.S.; analyzed the data: V.K., K.W., H.S.C., J.E., R.S, C.J.P.; wrote the paper: V.K., C.J.P.

