## Supplemental File for "Coordinated function of paired NLRs confers *Yr84*-mediated stripe rust resistance in wheat"

**Supplementary Text 1.** ***Conserved motifs in CNL and NL protein sequences***

The EDVID motif of the CC domain was detected by NLRexpress in both CNL and NL proteins. However, for NL, its similarity with VRELAYDAEDVID (Rairdan et al. 2008) was low (INALAERTGERLD) (Figure S5). Thus, we concluded that because the EDVID motif is likely missing, the NL protein lacks a functional CC domain. In contrast, the EDVID motif of the CNL protein is conserved and likely functional in mediating intramolecular interactions, as has been shown previously (Rairdan et al. 2008).

The NBD subdomain contains five conserved motifs, some or all of which were detected in CNL and NL protein sequences: VG, P-loop, RNBS-A, Walker-B and RNBS-B. The VG motif (bbGRE) (Martin et al. 2022) showed diversity in both CNL and NL proteins. The VG motif of NL was typical and was previously reported for CNL, TNL and RNL classes of NLRs as possessing the aa sequence VGIE/D. The motif of CNL (VGVE) (Figure S5) is not typical for previously described variances of VG motif (Martin et al. 2022).

The P-loop motif, responsible for what phosphate binding, (also termed Walker-A with consensus GbGGbGKTT) is highly conserved in CNL (GFGGLGKTT) but exhibits some variation in the NL protein sequence (GPVGVGKTA) (Figure 2A, S5, S6). Nevertheless, similar loss of conservation within this motif has been detected for some TNL proteins (Martin et al. 2022); thus, such variability is not conclusive for predicting its phosphate-binding functionality.

The motif RNBS-A was detected only for CNL. This highly hydrophobic motif plays a role in the formation of the nucleotide binding site. CNL protein holds conserved Phe in positions 0 and 5 of RNBS-A motif (Martin et al. 2022). In contrast, variation in the RNBS-A motif of NL made it impossible to detect (Figure S6). It has been noted in some publications that RNBS-A motif might be missing in some NLRs, such as these from TNL class (Steuernagel et al. 2015).

The Walker-B motif (KRFbbbbDDbW) is proposed to be involved in the coordination of the $Mg^{2+}$ ion required for ATPase activity of plant NLRs (Burdett et al. 2019). This motif is conserved in both, CNL (KRYLILIDDIW) and NL (RRYLIVIDDIW) (Figure 2A, S5, S6), suggesting their functionality.

The RNBS-B motif (KbbbTTR), involved in ligand binding, exhibits variability in the flanking regions for both CNL (RVVVTTR) and NL (RMVITTQ) (Figure S5, S6), as previously detected for many other NLR genes (Martin et al. 2022). However, this variability should not influence the functionality of this motif for CNL and NL, as the most critical amino acid, arginine in position 6, expected to actively participate in binding the nucleotide, remains conserved (Martin et al. 2022).

Within the ARC1 subdomain, two conserved motifs, RNBS-C and GLPL, were identified in both CNL and NL. In the RNBS-C motif, characterized by pattern LxxxExWxLF, both CNL (LDEEKSYNLF) and NL (LGELDSKNLL) (Figure 2A, S5, S6), exhibit high conservation of leucine aa at positions 0 and 8, which are functionally crucial as part of the binding site (Martin et al. 2022). The GLPL motif, essential for forming the binding site, is fully conserved in CNL, while NL displays only one amino acid change (GIPL), a common occurrence observed in analyses of large NLRs datasets (Martin et al. 2022).

Moving to the ARC2 subdomain, two more conserved motifs, RNBS-D and MHD, were annotated. The RNBS-D motif (CFbYCxLFP), as expected (Martin et al. 2022), exhibits high divergence for both CNL (CLLYMSMFP) and NL (LVLYMSMFP) (Figure 2A, S5, S6), with conservation observed at aa positions 3, 7 and 8 for both proteins. However, such variability is not critical for functionality, as this motif is primary involved in the NBS-LRR interface rather than ligan binding (Martin et al. 2022).

The MHD motifs in CNL and NL proteins show significant variation from the conserved structure. While CNL retains at least two critical amino acids (IHD), the MHD motif in NL was predicted to be completely different (VNY) (Figure 2A, S5, S6). The MHD motif, crucial for nucleotide-binding and regulation of subdomain interactions (Wang et al., 2020), is typically well-conserved. The detected drastic changes in the MHD domain of NL may signal its non-functionality.

**Supplementary Text 2.** ***Protein sequences of CNL-LRR and NL-LRR used for Co-Immunoprecipitation assays***

$CNL^{LRR}$ ~49 kD

MNFITIVGGHWQTPFPSYKVRRLSVQRSDEKETVQRMKLSHVRSLTAFESFKELHSCLLKFQIL
QVLDLESCRDLSSHRLDKICKMHQLKYLSLRRTDIGEIPSQIGMLEYLEVLDIRETKVRQLPKSV
KRLKQMTHLLTGNKSKRMAVTLTEEMTNMTALETLSGVGIYGSSRSNQVKTGIFKAASSGTYKE
VIQSLEKLTNLKKLSLYIIGMLEKANEESLLSAIEHLSSCSLKFLAIDDGFTGFLENSLSSSKAPPE
HLHTLELSGMLPKMPQWIVRLHSLGKLTLSFMSLKTETLILLAELPELFSLIFSIDAAKSNDPSVVK
ILYQNTLESGGEIIVPAGGFGKLKLLRLVAPVLPPLSFLGWAMPLLQRLELRFRISDGVHGLENLA
SLRQVFLTVSSKAPKAAEEIKGLASKIGSLPTVIVDEYNDQVEYLM

$NL^{LRR}$ ~ 52kD

MVARSNCNNILRRISLHQPDPDLQNFLNTRDLSHTRSLAISGTVDRVPLEKFVHLSVLDLEGWQ
RLKNKDLSLICNSGMFLLKYFSLRNTSISRLPPEIKKLLMLHTLVASHTQISELPSELCELGELESL
DLRSTQIRQLPEHIWKIPRLGTLLVGGDRVNCDLTITEVTEGMRFECYLRTLETVDLSECSACFV
EALACLFNLEKLSIRWSFGNCTDARLQKALCSCIKESRFFKSLTIHCGLGCQMDFLTSLPNPLPH
GSIEKFEVTGGSFVKIPQWFQGLQSLHSIEITVCRLEEGDVGILASLPSLVYLILGLRFVPEKVIVI
DRVGFPRLQKLSVCCRVPWLTFSSEAAMPSLTCLELKIGGGPRSEERIPSGFENLQNLWELVLS
YNEWYTMDSNVQTIVDAVTKGVAKHRYPNLITVIINGNPYKEGYVQEAREDAAKATRCQSATVR
HDPGNLQLFP

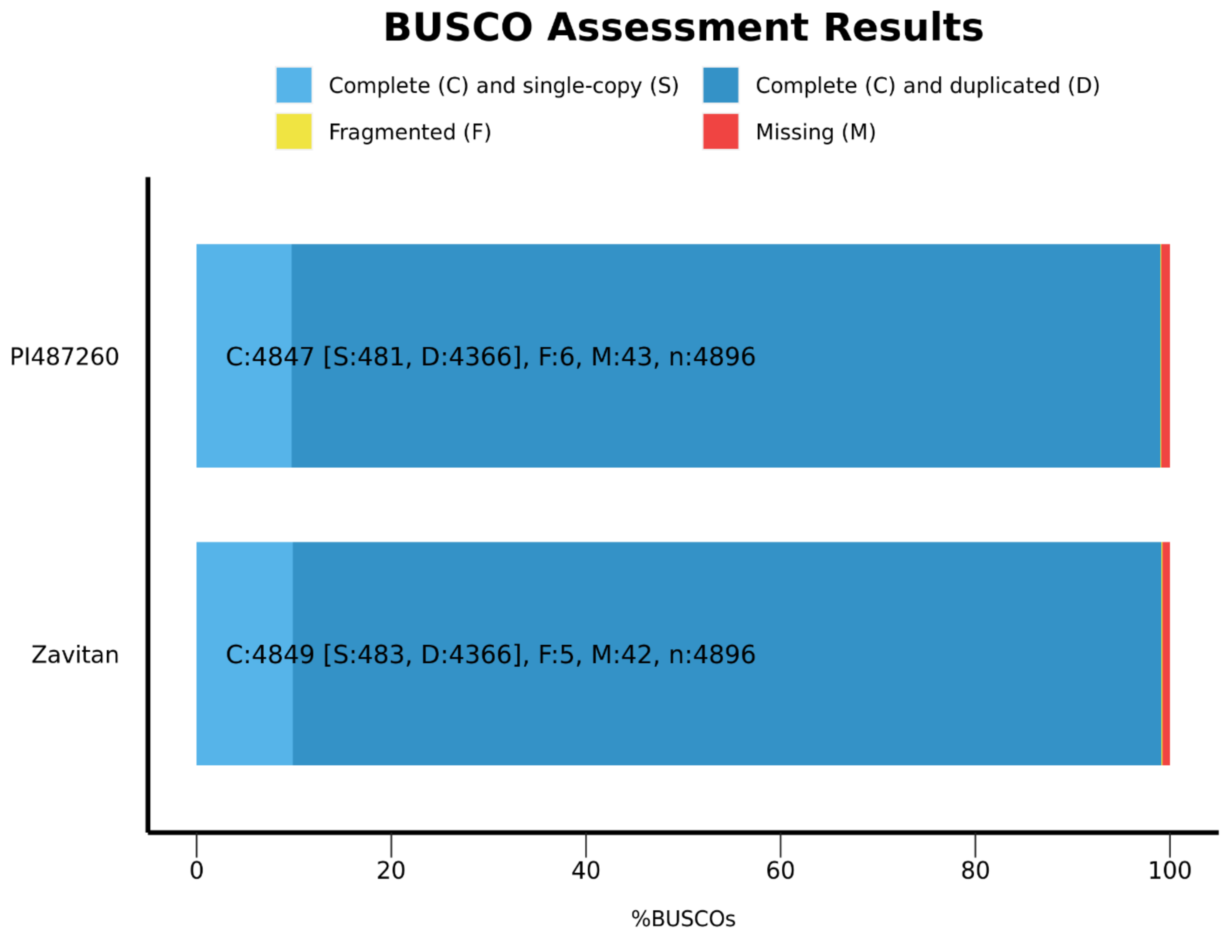


**Figure S1**. BUSCO analysis of the PI 487260 low-coverage CCS assembly compared with the wild emmer wheat "Zavitan" assembly (Avni et al. 2017).

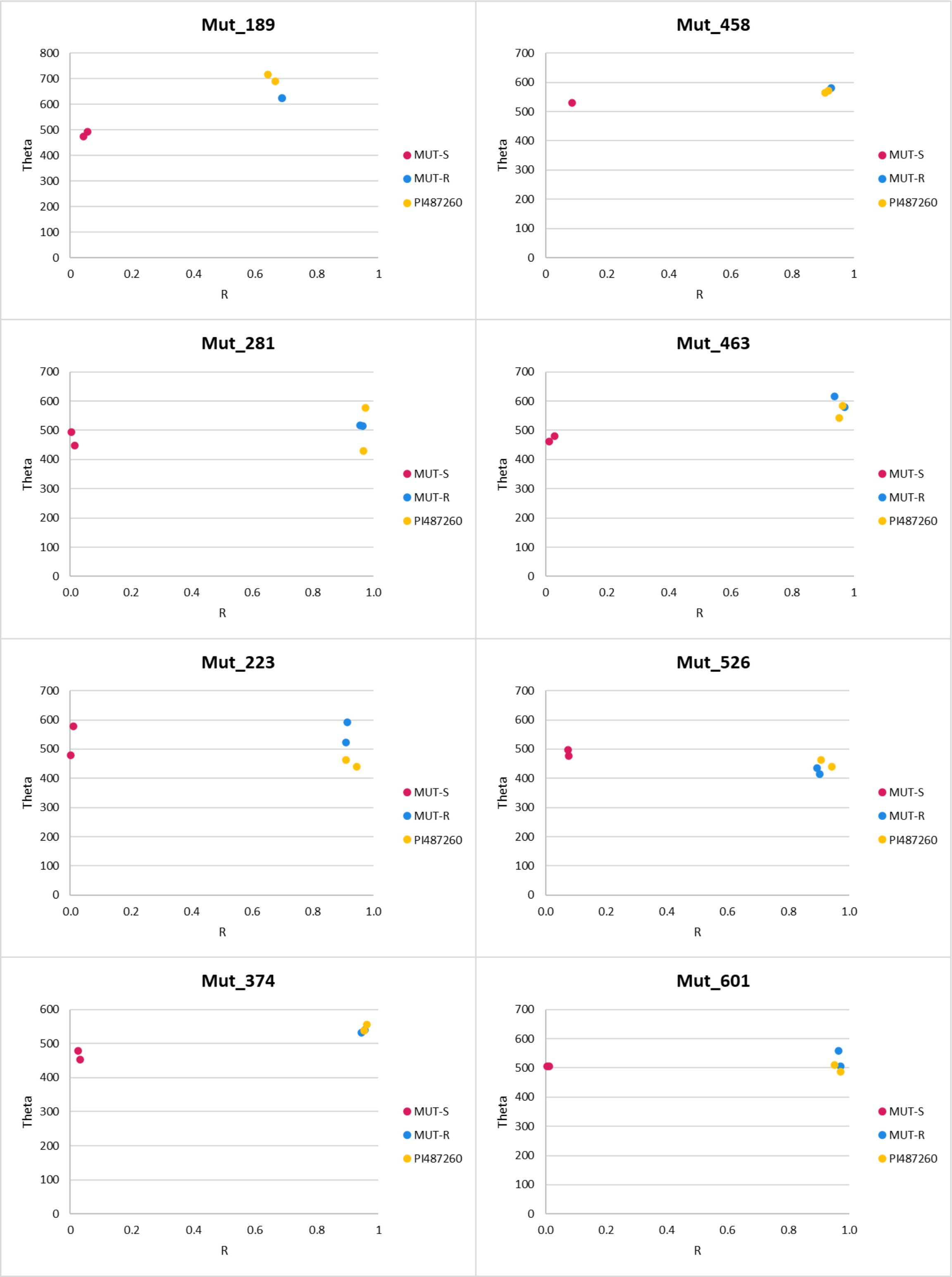


**Figure S2**. Theta graphs for KASP markers developed based on mutations detected in CNL and NL genes of susceptible mutants. All eight mutants show the presence of homozygous mutation (red), while all their corresponding sister lines (blue) and WT PI 487260 (yellow) show an absence of mutations.

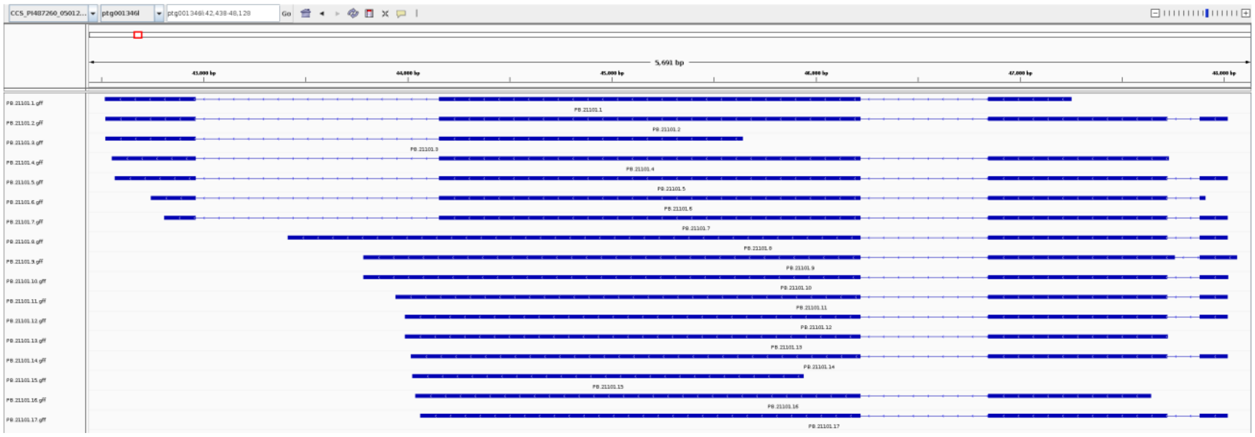

**Figure S3**. Seventeen CNL isoforms detected by Iso-Seq analysis. Translation of isoforms PB.21101.1, PB.21101.3, PB.21101.15, and PB.21101.16 results in truncated proteins, which are most likely not functional. All the remaining isoforms encode identical protein.

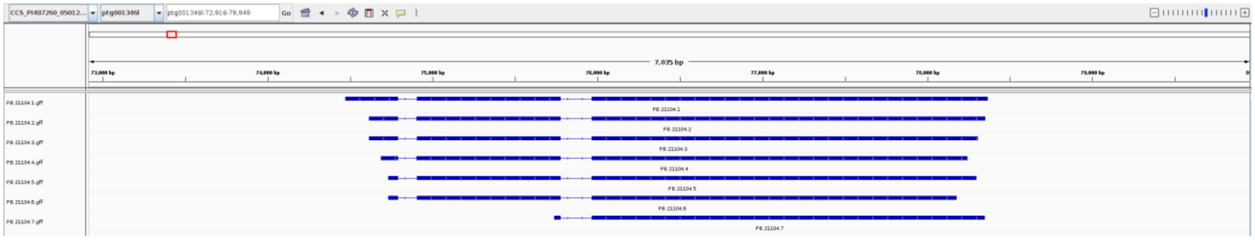

**Figure S4**. Seven NL isoforms detected by Iso-Seq analysis. The last two isoforms appear to be disrupted and most likely result in non-functional proteins: isoform PB.21104.7 is truncated, while isoform PB.21104.6 might carry a sequencing error, which change the ORF for protein prediction. The remaining isoforms encode almost identical proteins.

| Sequence | ResID | Module | Motif | | Proba(%) | Upstream | Motif sequence | Downstream |
|---|---|---|---|---|---|---|---|---|
| CNL | 73 | CCexpress | extEDVID | | 99.79 | WMKQV | REVSYDIEDCID | DVNHR |
| CNL | 168 | NBSexpress | VG | | 98.65 | TPPAE | LIGTK | QPVGV |
| CNL | 174 | NBSexpress | VG | | 99.24 | IGTKQ | PVGVE | DAIGE |
| CNL | 204 | NBSexpress | P-loop | | 100.00 | FLAID | GFGGLGKTT | LALEL |
| CNL | 225 | NBSexpress | RNSB-A | | 99.81 | QFGDG | FDCRAFVQVS | QKFDL |
| CNL | 282 | NBSexpress | Walker-B | | 97.81 | LQLKE | KRYLILID | DIWSV |
| CNL | 312 | NBSexpress | RNSB-B | | 98.46 | SSKGG | RVVVTTR | FRSVA |
| CNL | 339 | NBSexpress | RNSB-C | | 99.94 | YEHRP | LDEEKSYNLF | RQIIP |
| CNL | 372 | NBSexpress | GLPL | | 99.99 | LKKCG | GLPLV | IIVVA |
| CNL | 439 | NBSexpress | RNSB-D | | 98.75 | ADLKT | CLLYMSMFP | KGRSI |
| CNL | 508 | NBSexpress | MHD | | 99.65 | VKNYQ | IHD | MVLEY |
| CNL | 542 | LRRexpress | LxxLxL | | 99.06 | FPSYK | VRRLSV | QRSDE |
| CNL | 564 | LRRexpress | LxxLxL | | 93.24 | MKLSH | VRSLTA | FESFK |
| CNL | 586 | LRRexpress | LxxLxL | | 99.81 | LKFQI | LQVLDL | ESCRD |
| CNL | 611 | LRRexpress | LxxLxL | | 99.95 | CKMHQ | LKYLSL | RRTDI |
| CNL | 634 | LRRexpress | LxxLxL | | 99.96 | GMLEY | LEVLDI | RETKV |
| CNL | 657 | LRRexpress | LxxLxL | | 99.58 | KRLKQ | MTHLLT | GNKSK |
| CNL | 686 | LRRexpress | LxxLxL | | 28.91 | TALET | LSGVGI | YGSSR |
| CNL | 726 | LRRexpress | LxxLxL | | 99.64 | EKLTN | LKKLSL | YIIGM |
| CNL | 752 | LRRexpress | LxxLxL | | 38.22 | SAIEH | LSSCSL | KFLAI |
| CNL | 757 | LRRexpress | LxxLxL | | 43.60 | LSSCS | LKFLAI | DDGFT |
| CNL | 784 | LRRexpress | LxxLxL | | 99.93 | APPEH | LHTLEL | SGMLP |
| CNL | 806 | LRRexpress | LxxLxL | | 99.91 | VRLHS | LGKLTL | SFMSL |
| CNL | 830 | LRRexpress | LxxLxL | | 98.42 | AELPE | LFSLIF | SIDAA |
| CNL | 872 | LRRexpress | LxxLxL | | 92.10 | GGFGK | LKLLRL | VAPVL |
| CNL | 895 | LRRexpress | LxxLxL | | 99.92 | WAMPL | LQRLEL | RFRIS |
| CNL | 917 | LRRexpress | LxxLxL | | 98.25 | ENLAS | LRQVFL | TVSSK |

**Figure S5**. Conserved motifs identified in CNL protein using NLRexpress. ResID – residue ID where the predicted motif starts, Proba (%) – inferred probability to start the given motif, Upstream – five residues upstream the motif, Downstream – five residues downstream the motif end.

| Sequence | ResID | Module | Motif | | Proba(%) | Upstream | Motif sequence | Downstream |
|---|---|---|---|---|---|---|---|---|
| NL | 102 | CCexpress | extEDVID | | 96.12 | WLYGI | NALAERTGERLD | KREQS |
| NL | 184 | NBSexpress | VG | | 99.24 | ESSMP | LVGIE | STLAN |
| NL | 215 | NBSexpress | P-loop | | 100.00 | IISMV | GPVGVGKTA | IAMDV |
| NL | 293 | NBSexpress | Walker-B | | 97.63 | QCLQD | RRYLIVID | DIWEI |
| NL | 323 | NBSexpress | RNSB-B | | 98.46 | NSRGS | RMVITTQ | IRSIA |
| NL | 350 | NBSexpress | RNSB-C | | 99.94 | HEVKS | LGELDSKNLL | FRKAF |
| NL | 388 | NBSexpress | GLPL | | 99.95 | LRRCE | GIPLF | INGMA |
| NL | 443 | NBSexpress | RNSB-D | | 98.75 | YQMKV | LVLYMSMFP | DGYIF |
| NL | 512 | NBSexpress | MHD | | 99.65 | DCHYK | VNY | FMLQF |
| NL | 547 | LRRexpress | LxxLxL | | 98.70 | NCNNI | LRRISL | HQPDP |
| NL | 571 | LRRexpress | LxxLxL | | 74.65 | RDLSH | TRSLAI | SGTVD |
| NL | 591 | LRRexpress | LxxLxL | | 99.57 | EKFVH | LSVLDL | EGWQR |
| NL | 618 | LRRexpress | LxxLxL | | 99.88 | SGMFL | LKYFSL | RNTSI |
| NL | 641 | LRRexpress | LxxLxL | | 99.94 | KKLLM | LHTLVA | SHTQI |
| NL | 664 | LRRexpress | LxxLxL | | 99.96 | CELGE | LESLDL | RSTQI |
| NL | 687 | LRRexpress | LxxLxL | | 99.96 | WKIPR | LGTLLV | GGDRV |
| NL | 719 | LRRexpress | LxxLxL | | 88.70 | CYLRT | LETVDL | SECSA |
| NL | 741 | LRRexpress | LxxLxL | | 99.48 | ACLFN | LEKLSI | RWSFG |
| NL | 772 | LRRexpress | LxxLxL | | 96.69 | KESRF | FKSLTI | HCGLG |
| NL | 800 | LRRexpress | LxxLxL | | 99.91 | LPHGS | IEKFEV | TGGSF |
| NL | 823 | LRRexpress | LxxLxL | | 99.97 | QGLQS | LHSIEI | TVCRL |
| NL | 847 | LRRexpress | LxxLxL | | 97.67 | ASLPS | LVYLIL | GLRFV |
| NL | 872 | LRRexpress | LxxLxL | | 99.15 | VGFPR | LQKLSV | CCRVP |
| NL | 895 | LRRexpress | LxxLxL | | 99.48 | AAMPS | LTCLEL | KIGGG |
| NL | 922 | LRRexpress | LxxLxL | | 96.60 | ENLQN | LWELVL | SYNEW |
| NL | 958 | LRRexpress | LxxLxL | | 68.27 | HRYPN | LITVII | NGNPY |

**Figure S6**. Conserved motifs found in NL protein using NLRexpress. ResID – residue ID where the predicted motif starts, Proba (%) – inferred probability to start the given motif, Upstream – five residues upstream the motif, Downstream – five residues downstream the motif end.

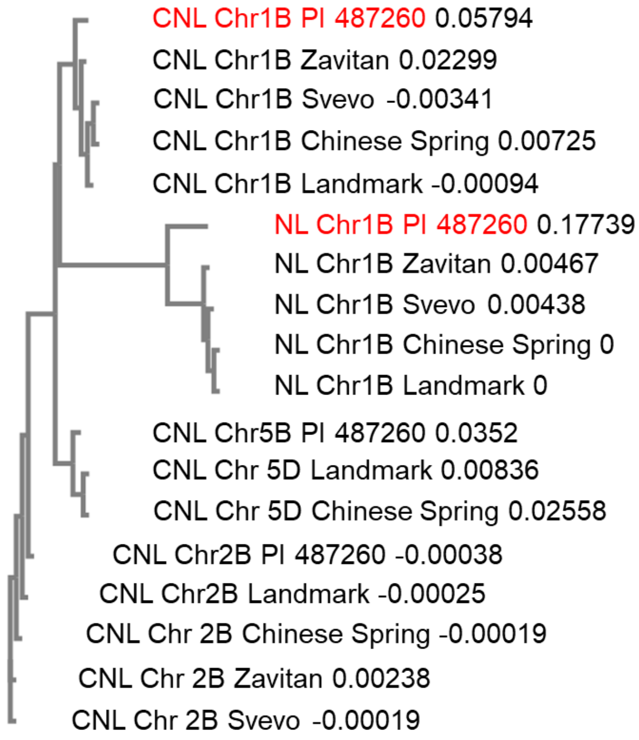


**Figure S7**. Phylogenetic tree of CNL and NL copies found across reference genomes of hexaploid wheat (CDC Landmark, Chinese Spring) and tetraploid wheat (Zavitan, Svevo, and PI 487260). The tree is based on CDS sequences and was visualized using EMBL-EBI Simple Phylogeny online tool. CNL and NL functional alleles are highlighted in red.

**Table S1**. Haplotypes of 56 $F_2$ recombinants identified after screening of 1039 $F_2$ plants. Green – homozygous PI 487260 allele, yellow – homozygous Kronos allele, blue – heterozygous. Lines selected for development of homozygous $F_{2:3}$ recombinants are marked in bold.

| Sample | *usw310* | *usw312* | *usw313* | Pst IT | *usw314* | *usw315* | *usw316* | *usw317* | *usw318* |
|---|---|---|---|---|---|---|---|---|---|
| PI487260 | | | | 2 | | | | | |
| Kronos | | | | 9 | | | | | |
| **KSY_R1_348** | | | | 4 | | | | | |
| **KSY_R2_166** | | | | 4 | | | | | |
| KSY_R1_89 | | | | 3 | | | | | |
| KSY_R2_49 | | | | 5 | | | | | |
| KSY_R2_111 | | | | 4 | | | | | |
| KSY_R2_306 | | | | 3 | | | | | |
| KSY_R2_316 | | | | 6 | | | | | |
| KSY_R3_75 | | | | 2 | | | | | |
| KSY_R3_76 | | | | 3 | | | | | |
| KSY_R3_84 | | | | 6 | | | | | |
| KSY_R3_116 | | | | 4 | | | | | |
| KSY_R3_163 | | | | 4 | | | | | |
| KSY_R3_168 | | | | 6 | | | | | |
| KSY_R3_226 | | | | 1 | | | | | |
| KSY_R3_250 | | | | 6 | | | | | |
| KSY_R3_275 | | | | 4 | | | | | |
| **KSY_R1_256** | | | | 3 | | | | | |
| **KSY_R1_296** | | | | 1 | | | | | |
| KSY_R3_148 | | | | 1 | | | | | |
| KSY_R1_141 | | | | - | | | | | |
| KSY_R1_226 | | | | 1 | | | | | |
| KSY_R1_239 | | | | 2 | | | | | |
| KSY_R2_81 | | | | 0 | | | | | |
| **KSY_R1_118** | | | | 9 | | | | | |
| **KSY_R2_326** | | | | 9 | | | | | |
| KSY_R2_167 | | | | 9 | | | | | |
| KSY_R2_193 | | | | 9 | | | | | |
| KSY_R2_220 | | | | 9 | | | | | |
| KSY_R2_242 | | | | 9 | | | | | |
| KSY_R2_307 | | | | 9 | | | | | |
| KSY_R3_32 | | | | 9 | | | | | |
| KSY_R3_34 | | | | 9 | | | | | |
| KSY_R1_359 | | | | 9 | | | | | |
| KSY_R3_233 | | | | 9 | | | | | |
| KSY_R3_279 | | | | 9 | | | | | |
| **KSY_R1_298** | | | | 5 | | | | - | |
| **KSY_R2_70** | | | | 4 | | | | | |
| KSY_R2_255 | | | | 3 | | | | | |
| KSY_R2_312 | | | | 6 | | | | | |
| **KSY_R1_64** | | | | 2 | | | | | |
| **KSY_R1_152** | | | | 6 | | | | | |
| **KSY_R3_241** | | | | 4 | | | | | |
| **KSY_R2_236** | | | | 2 | | | | | |
| **KSY_R1_315** | | | | 6 | | | | | |
| **KSY_R1_356** | | | | 4 | | | | | |
| **KSY_R3_334** | | | | 6 | | | | | |
| KSY_R2_207 | | | | 3 | | | | | |
| **KSY_R2_289** | | | | 7 | | | | | |
| **KSY_R1_319** | | | | 9 | | | | | |
| **KSY_R3_42** | | | | 9 | | | | | |
| KSY_R1_245 | | | | 9 | | | | | |
| **KSY_R2_42** | | | | 1 | | | | | |
| **KSY_R2_233** | | | | 1 | | | | | |
| KSY_R3_13 | | | | 1 | | | | | |
| KSY_R3_52 | | | | 1 | | | | | |
| KSY_R3_71 | | | | 1 | | | | | |

**Table S2**. Haplotypes of $F_{2:3}$ homozygous recombinants from Kronos×PI 487260 population. Green – homozygous PI 487260 allele, yellow – homozygous Kronos allele. Bold box outlines *Yr84* genetic region.

| Sample | *usw310* | *usw312* | *usw313* | ***Yr84*** | *usw314* | *usw315* | *usw316* | *usw317* |
|---|---|---|---|---|---|---|---|---|
| PI487260 | | | | 2 | | | | |
| Kronos | | | | 9 | | | | |
| KSY_R1_256_9 | | | | 1 | | | | |
| KSY_R1_296_2 | | | | 1 | | | | |
| KSY_R1_298_7 | | | | 1 | | | | |
| KSY_R2_70_3 | | | | 1 | | | | |
| KSY_R1-64-3 | | | | 1 | | | | |
| KSY_R1_152_10 | | | | 7 | | | | |
| KSY_R3_241_5 | | | | 9 | | | | |
| KSY_R2_289_9 | | | | 9 | | | | |
| KSY_R1_319_1 | | | | 9 | | | | |
| KSY_R3_42_4 | | | | 9 | | | | |
| KSY_R1_348_1 | | | | 9 | | | | |
| KSY_R2_166_3 | | | | 9 | | | | |
| KSY_R1_118_2 | | | | 9 | | | | |
| KSY_R2_326_7 | | | | 9 | | | | |
| KSY_R2_236_4 | | | | 1 | | | | |
| KSY_R1_315_10 | | | | 1 | | | | |
| KSY_R1_356_7 | | | | 1 | | | | |
| KSY_R3_334_3 | | | | 1 | | | | |
| KSY_R2_42_3 | | | | 1 | | | | |
| KSY_R2_233_1 | | | | 1 | | | | |

**Table S3**. Candidate genes (and associated annotations) in the *Yr84* genomic region (*usw313-usw316*) of the Zaviatan wild emmer wheat assembly.

| Zavitan gene | Start | End | Contig | Start | End | Annotation |
|---|---|---|---|---|---|---|
| ***TRIDC1BG002180.1.mrna1*** | ***11032887*** | ***11035943*** | ***ptg001346l*** | ***42520*** | ***48019*** | ***Disease resistance protein*** |
| ***TRIDC1BG002190.1.mrna1*** | ***11047307*** | ***11049713*** | ***ptg001346l*** | ***74472*** | ***78366*** | ***Disease resistance protein (NBS-LRR class) family*** |
| **TRIDC1BG002210.1.mrna1** | **11095915** | **11096246** | **ptg001346l** | **165910** | **166251** | **Defensin** |
| TRIDC1BG002200.2.mrna1 | 11178525 | 11179885 | ptg001346l | 225574 | 226923 | Glutathione S-transferase family protein |
| TRIDC1BG002220.3.mrna1 | 11188633 | 11195858 | ptg001346l | 209182 | 216017 | Methionine S-methyltransferase |
| TRIDC1BG002230.1.mrna1 | 11199032 | 11201334 | ptg001346l | 196760 | 199359 | U-box domain protein |
| **TRIDC1BG002260.1.mrna1** | **11205370** | **11205870** | | | | **Serine/threonine-protein kinase CTR1** |
| **TRIDC1BG002270.1.mrna1** | **11211587** | **11213098** | **ptg001346l** | **595315** | **596820** | **Disease resistance protein RGA2** |
| TRIDC1BG002280.1.mrna1 | 11228824 | 11230656 | ptg001346l | 612860 | 614692 | Biosynthetic arginine decarboxylase |
| TRIDC1BG002290.2.mrna1 | 11246308 | 11248683 | ptg001346l | 628941 | 630392 | C2 calcium/lipid-binding plant phosphoribosyl transferase family protein |
| **TRIDC1BG002300.1.mrna1** | **11270911** | **11271084** | **ptg001346l** | **655101** | **655263** | **Defensin** |
| **TRIDC1BG002310.1.mrna1** | **11275738** | **11276083** | **ptg001346l** | **660110** | **660397** | **Defensin** |
| **TRIDC1BG002340.1.mrna1** | **11425210** | **11427732** | **ptg000814l** | **370563** | **371321** | **Serine/threonine-protein kinase** |
| TRIDC1BG002350.2.mrna1 | 11473877 | 11474122 | ptg000814l | 209912 | 210154 | Glutathione S-transferase |
| TRIDC1BG002370.8.mrna1 | 11476963 | 11481821 | ptg000814l | 213207 | 218042 | Tubulin-specific chaperone cofactor E-like protein |
| TRIDC1BG002380.1.mrna1 | 11483667 | 11487378 | ptg000814l | 221175 | 225228 | Rab GTPase family 1 |
| **TRIDC1BG002390.2.mrna1** | **11495931** | **11496689** | **ptg000814l** | **233680** | **233855** | **Disease resistance protein (NBS-LRR class) family** |
| **TRIDC1BG002400.1.mrna1** | **11497198** | **11497887** | **ptg000814l** | **234525** | **234762** | **Disease resistance protein RPM1** |
| TRIDC1BG002410.2.mrna1 | 11501055 | 11502841 | ptg000814l | 237303 | 239096 | Ras-related protein Rab-25 |
| TRIDC1BG002420.2.mrna1 | 11571371 | 11572619 | ptg000814l | 337550 | 338798 | 12-oxophytodienoate reductase-like protein |
| TRIDC1BG002430.1.mrna1 | 11577115 | 11578744 | ptg000814l | 352234 | 353861 | 12-oxophytodienoate reductase-like protein |
| **TRIDC1BG002460.2.mrna1** | **11941709** | **11943554** | **ptg000814l** | **560737** | **561634** | **Protein kinase family protein** |

Genes **in bold** were annotated as putative disease resistance genes.
Genes in blue highlight are the *Yr84* paired NLRs.

**Table S4:** Assembly statistics for the CCS assembly of PI 487260.

| **Parameter** | |
|---|---|
| No. contigs | 4615 |
| Largest contig, Mbp | 54.5 |
| Total length, Mbp | 10460.1 |
| GC (%) | 46.1 |
| $N_{50}$, Mbp | 6.7 |
| $N_{75}$, Mbp | 3.5 |
| $L_{50}$ | 440 |
| $L_{75}$ | 979 |

**Table S5.** Summary of paired-end raw read data generated for each of the susceptible mutants. De-multiplexed reads were aligned to PI 487260 assembly.

| Sample | Pool | Index | PE reads | coverage |
|---|---|---|---|---|
| Mut_223 | PI487260mut_susceptible-pool-1 | UDP0153 | 736,698,377 | 12 |
| Mut_374 | PI487260mut_susceptible-pool-1 | UDP0155 | 614,494,624 | 10 |
| Mut_189 | PI487260mut_susceptible-pool-1 | UDP0157 | 791,863,544 | 13 |
| Mut_281 | PI487260mut_susceptible-pool-1 | UDP0158 | 755,084,399 | 12 |
| Mut_458 | PI487260mut_susceptible-pool-2 | UDP0161 | 330,022,753 | 5 |
| Mut_463 | PI487260mut_susceptible-pool-2 | UDP0162 | 588,889,449 | 10 |
| Mut_479 | PI487260mut_susceptible-pool-2 | UDP0163 | 681,345,724 | 11 |
| Mut_526 | PI487260mut_susceptible-pool-2 | UDP0164 | 761,832,443 | 12 |
| Mut_601 | PI487260mut_susceptible-pool-2 | UDP0165 | 661,081,544 | 11 |

**Table S6.** Primers used in the current study.

| Experiment | Primer name | Primer sequence* |
|---|---|---|
| CNL/NL amplicon sequencing | CNL_amp | F: CACGAGCAGCAGTGGTAGCCT<br>R: TCAGCCATCCATAGCAGTTGA |
| | NL_amp | F: CTGCGTGTACTAAGAGACTAG<br>R: TTACGCGCAGTGTTCGCTCTA |
| Expression studies | CNL_exp | F: CTTGTTGTCACCACTACTCTG<br>R: CAATTAAAAGAGAAGAGGTAC |
| | NL_exp | F: GTACTGCGCTACGGCTAAGTT<br>R: TCAATCACGATGAGGTACCTC |
| Mutations in *Yr84* susceptible mutants | Mut_189 | HF: GAAGGTCGGAGTCAACGGATTTGGCATGTTACCAAAAATGCT<br>FF: GAAGGTGACCAAGTTCATGCTTGGCATGTTACCAAAAATGCC<br>R: CTCAGGCAGTTCAGCAAGAAG |
| | Mut_223 | HF: GAAGGTCGGAGTCAACGGATTACGTGAGGAAGGCGACAACTA<br>FF: GAAGGTGACCAAGTTCATGCTACGTGAGGAAGGCGACAACTT<br>R: CATCAACGGATGCAAGCATTG |
| | Mut_281 | HF: GAAGGTCGGAGTCAACGGATTATTGTACATGAGCATGTTTCT<br>FF: GAAGGTGACCAAGTTCATGCTATTGTACATGAGCATGTTTCC<br>R: CGAATCAAGTTCCGGACAATC |
| | Mut_374 | HF: GAAGGTCGGAGTCAACGGATTTTACCAAAAATGCCCCAGTGA<br>FF: GAAGGTGACCAAGTTCATGCTTTACCAAAAATGCCCCAGTGG<br>R: CTCAGGCAGTTCAGCAAGAAG |
| | Mut_458 | HF: GAAGGTCGGAGTCAACGGATTTTTCATCACAATAGTGGGTGA<br>FF: GAAGGTGACCAAGTTCATGCTTTTCATCACAATAGTGGGTGG<br>R: TTCTCTGGACAGACAACCTGC |
| | Mut_463 | HF: GAAGGTCGGAGTCAACGGATTATGAATTTTCAAGAAAGCCGA<br>FF: GAAGGTGACCAAGTTCATGCTATGAATTTTCAAGAAAGCCGG<br>R: GCTGCTCCCTCAAGTTTCTGG |
| | Mut_526 | HF: GAAGGTCGGAGTCAACGGATTTCCTGTTGTTTTTGGTGCAAT<br>FF: GAAGGTGACCAAGTTCATGCTTCCTGTTGTTTTTGGTGCAAC<br>R: ATTAGGCAAGTTTGCAGCAAC |
| | Mut_601 | HF: GAAGGTCGGAGTCAACGGATTCTACTCTGCCACCCTTGGAAT<br>FF: GAAGGTGACCAAGTTCATGCTCTACTCTGCCACCCTTGGAAC<br>R: ATGTGTAGGTACCTTATCTTG |
| Y2H | CNL_pGADT7 AD | F: GGAGGCCAGTGAATTCATGGACCTCGTCGTCGGC<br>R: CGAGCTCGATGGATCCCATCAAATACTCTACTTGATCATTG |
| | CNL_pGBKT7 | F: CATGGAGGCCGAATTCATGGACCTCGTCGTCGGC<br>R: GCAGGTCGACGGATCCCATCAAATACTCTACTTGATCATTG |
| | NL_pGADT7 AD | F: GGAGGCCAGTGAATTCATGGCTGAGGATGGGGGAG<br>R: CGAGCTCGATGGATCCAGGAAATAACTGCAGATTGCCAGG |
| | NL_pGBKT | F: CATGGAGGCCGAATTCATGGCTGAGGATGGGGGAG<br>R: GCAGGTCGACGGATCCAGGAAATAACTGCAGATTGCCAGG |
| PI 487260xPI 471699 | *usw329* | HF: GAAGGTCGGAGTCAACGGATTCGCTAGCCACTCTGTCTAAGTAC<br>FF: GAAGGTGACCAAGTTCATGCTCGCTAGCCACTCTGTCTAAGTAT<br>R: CGCGTCCACTTGTTTATACCA |
| | *usw330* | HF: GAAGGTCGGAGTCAACGGATTTCCCCACTTCCAACAAGTCC<br>FF: GAAGGTGACCAAGTTCATGCTTCCCCACTTCCAACAAGTCT<br>R: CCCGGAAATTACCTCAGCCA |

*F – forward primer, R – reverse primer, HF – HEX forward primer, FF – FAM forward primer

**Table S7:** PI 487260 Iso-Seq sequencing statistics

| **Parameter** | **Count** |
|---|---|
| Total sequencing data (bp) | 9,001,476,096 |
| Mean length of reads (bp) | 2,329 |
| Total number of Hifi reads | 3,626,159 |
| Full-Length Non-Chimeric Reads | 3,618,635 |
| Full-Length Non-Chimeric Reads with Poly-A Tail | 3,615,628 |
| Number of mapped unique isoforms | 155,477 |
| Number of mapped unique loci | 29,745 |

**Table S8.** CNL and NL haplotypes based on amplicon sequencing of 38 WEW accession. Phenotypic response to *Pst* race W001 inoculation and presence of *Yr15* (*WTK1*) based on functional KASP markers (Klymiuk et al., 2019) are indicated. Only one genotype, PI 471699, showed a resistance response in the absence of *Yr15* gene and possessed unique haplotypes for both genes. However, the PI 487260 x PI 471699 $F_2$ mapping population showed presence of two dominant R genes, and the CNL and NL alleles of PI 471699 did not confer resistance (see main text and Supplementary Table S11 for details).

| Geographic location | Accession | CNL | NL | *usw314* allele | *Pst W001* phenotype (IT) | *Yr15 (WTK1)* |
|---|---|---|---|---|---|---|
| Syria, As Suwaydā' | **PI 487260** | Hap1 | Hap1 | A | 2 | - |
| Iran | **IG113301** | Hap5 | Hap2 | A | 9 | - |
| Iraq | **IG131233** | Hap6 | Hap3 | A | 9 | - |
| Jordan | **IG139169** | Hap2 | Hap5 | A | 9 | - |
| Jordan | **IG140059** | Hap2 | Hap5 | A | 9 | - |
| Jordan | **IG46057** | Hap2 | Hap12 | A | 9 | - |
| Jordan | **IG46374** | Hap4 | Hap6 | A | 9 | - |
| Jordan | **IG46387** | Hap3 | Hap4 | A | 7 | - |
| Israel, Maghar | **PI 415148** | Hap4 | Hap7 | A | 6 | - |
| Israel, Eliad | **PI 471699** | Hap10 | Hap15 | A | 3 | - |
| Israel, Beit Rimon | **PI 471720** | Hap7 | Hap8 | A | 9 | - |
| Israel, Safed | **PI 471725** | Hap7 | Hap8 | A | 9 | - |
| Israel, T.A.P. Yoav camp | **PI 471739** | Hap7 | Hap8 | A | 2 | + |
| Israel, Jerusalem | **PI 471753** | Hap4 | Hap9 | A | 9 | - |
| Israel, Beit Keshet | **PI 471766** | Hap2 | Hap13 | A | 9 | - |
| Israel, Dalia | **PI 471814** | Hap2 | Hap9 | A | 9 | - |
| Israel, Kadarim | **PI 478682** | Hap4 | Hap9 | A | 8 | - |
| Israel, Tel Susita | **PI 478684** | Hap7 | Hap8 | A | 9 | - |
| Israel, Mt. Moreh | **PI 478686** | Hap8 | Hap8 | A | 8 | - |
| Israel, Eilabun | **PI 478717** | Hap4 | Hap9 | A | 9 | - |
| Israel, Tekoa | **PI 478718** | Hap4 | Hap9 | A | 9 | - |
| Israel, Goren | **PI 478723** | Hap4 | Hap9 | A | 9 | - |
| Israel, Ari'el | **PI 478737** | N/A | Hap14 | A | 7 | - |
| Israel, Kfar Hittim | **PI 481485** | Hap4 | Hap9 | A | 9 | - |
| Israel, Har Savyon | **PI 481524** | Hap4 | Hap9 | A | 9 | - |
| Israel, Amirim | **TD011913** | Hap4 | Hap9 | A | 9 | - |
| Israel, Nahef | **TD011993** | Hap4 | Hap9 | A | 9 | - |
| Israel, Zavitan | **Zavitan** | N/A | Hap9 | B | 9 | - |
| Syria, Rīf Dimashq | **PI 487256** | Hap9 | Hap8 | A | 9 | - |
| Syria, Rīf Dimashq | **PI 487257** | Hap6 | Hap10 | A | 9 | - |
| Syria, Rīf Dimashq | **PI 487259** | Hap6 | Hap10 | A | 8 | - |
| Lebanon, Al Biqāʻ | **PI 428128** | Hap2 | Hap12 | A | 2 | + |
| Lebanon, Béqaa | **PI 428136** | Hap2 | Hap12 | A | 3 | + |
| Lebanon, Béqaa | **PI 538701** | Hap2 | N/A | A | 2 | + |

| | | | | | | |
|---|---|---|---|---|---|---|
| Lebanon, Béqaa | **PI 538710** | Hap2 | Hap12 | A | 2 | + |
| Turkey, Şırnak | **PI 560697** | N/A | Hap12 | H | 9 | - |
| Turkey, Şanlıurfa | **PI 654325** | Hap5 | Hap11 | A | 9 | - |
| Turkey, Gaziantep | **PI 654334** | Hap5 | Hap11 | A | 8 | - |
| Turkey, Kilis | **PI 654336** | Hap5 | Hap11 | A | 9 | - |
| Turkey, Kahramanmaraş | **PI 656872** | N/A | Hap9 | H | 9 | - |

**Table S9.** Exon CNL haplotypes found in wild emmer wheat natural populations.

| **Haplotype** | **Nucleotide position(s) relative to the start codon of the CNL gene in gDNA (only exons are presented)** | | | | | | | | **N of genotypes** |
|---|---|---|---|---|---|---|---|---|---|
| | **87-107** | **228** | **248** | **279** | **2941** | **3192** | **3221** | **3290** | |
| **Hap1** | CATCCAGGGCGTCCATGACGA | C | T | G | T | T | G | G | 1 |
| **Hap2** | CATCCAGGGCGTCCATGACGA | C | T | G | T | T | **A** | G | 9 |
| **Hap3** | CATCCAGGGCGTCCATGACGA | **T** | T | G | T | T | **A** | G | 1 |
| **Hap4** | deletion | C | **C** | G | **A** | **C** | **A** | G | 11 |
| **Hap5** | CATCCAGGGCGTCCATGACGA | C | T | G | **A** | **C** | **A** | G | 4 |
| **Hap6** | CATCCAGGGCGTCCATGACGA | C | T | G | T | T | **A** | G | 3 |
| **Hap7** | CATCCAGGGCGTCCATGACGA | C | T | G | T | T | **A** | **A** | 4 |
| **Hap8** | CATCCAGGGCGTCCATGACGA | C | T | G | T | T | **A** | **A** | 1 |
| **Hap9** | CATCCAGGGCGTCCATGACGA | C | T | G | **C** | T | **A** | **A** | 1 |
| **Hap10** | CATCCAGGGCGTCCATGACGA | C | T | **A** | T | T | **A** | G | 1 |

**Table S10.** Exon NL haplotypes found in wild emmer wheat natural populations.

| **Haplotype** | **Nucleotide position(s) relative to the start codon of the NL gene in gDNA (only exons are presented)** | | | | | | | | | **N of genotypes** |
|---|---|---|---|---|---|---|---|---|---|---|
| | **335** | **441** | **636** | **1232** | **1681** | **2021** | **2182** | **2374** | **3182** | |
| **Hap1** | G | G | T | A | A | C | C | A | G | 1 |
| **Hap2** | G | G | **A** | A | A | C | C | A | G | 1 |
| **Hap3** | G | G | T | **T** | A | C | C | A | **A** | 1 |
| **Hap4** | G | G | T | **T** | A | C | C | A | **A** | 1 |
| **Hap5** | G | **A** | T | A | A | **G** | C | **G** | G | 2 |
| **Hap6** | G | G | **A** | A | **G** | C | C | A | G | 1 |
| **Hap7** | G | G | **A** | A | A | C | C | A | G | 1 |
| **Hap8** | **C** | G | T | A | A | C | C | A | G | 6 |
| **Hap9** | G | G | **A** | A | A | C | C | A | G | 12 |
| **Hap10** | G | G | T | **T** | A | C | C | A | **A** | 2 |
| **Hap11** | G | G | **A** | A | A | C | C | A | G | 3 |
| **Hap12** | G | G | T | A | A | C | C | A | G | 5 |
| **Hap13** | G | **A** | T | A | A | C | C | A | G | 1 |
| **Hap14** | G | G | T | **T** | A | C | **A** | A | G | 1 |
| **Hap15** | G | G | T | A | A | C | **T** | A | G | 1 |

**Table S11:** *Pst* phenotyping and genotyping of an $F_2$ population derived from the cross PI 487160 x PI 471699 $F_2$ population. Homozygous PI 487260 allele marked as 1, homozygous PI 471699 allele – 0, heterozygous allele – H. *Pst* infection type (IT) on a 0-9 scale. Presence of PI 471699 CNL/NL alleles did not always confer the resistance; thus, they are non-functional.

| Genotypes | *usw329* (CNL) | *usw330* (NL) | *Pst* IT |
|---|---|---|---|
| PI 487260 | 1 | 1 | 1 |
| PI 471699 | 0 | 0 | 2 |
| PIxPI_142 | 0 | 0 | 0 |
| PIxPI_108 | 1 | 1 | 0 |
| PIxPI_167 | 1 | 1 | 0 |
| PIxPI_169 | 1 | 1 | 0 |
| PIxPI_27 | 0 | 0 | 1 |
| PIxPI_174 | 0 | 0 | 1 |
| PIxPI_74 | 0 | H | 1 |
| PIxPI_4 | 1 | 1 | 1 |
| PIxPI_5 | 1 | 1 | 1 |
| PIxPI_6 | 1 | 1 | 1 |
| PIxPI_8 | 1 | 1 | 1 |
| PIxPI_9 | 1 | 1 | 1 |
| PIxPI_10 | 1 | 1 | 1 |
| PIxPI_16 | 1 | 1 | 1 |
| PIxPI_21 | 1 | 1 | 1 |
| PIxPI_32 | 1 | 1 | 1 |
| PIxPI_49 | 1 | 1 | 1 |
| PIxPI_59 | 1 | 1 | 1 |
| PIxPI_68 | 1 | 1 | 1 |
| PIxPI_72 | 1 | 1 | 1 |
| PIxPI_75 | 1 | 1 | 1 |
| PIxPI_81 | 1 | 1 | 1 |
| PIxPI_86 | 1 | 1 | 1 |
| PIxPI_92 | 1 | 1 | 1 |
| PIxPI_101 | 1 | 1 | 1 |
| PIxPI_102 | 1 | 1 | 1 |
| PIxPI_110 | 1 | 1 | 1 |
| PIxPI_130 | 1 | 1 | 1 |
| PIxPI_135 | 1 | 1 | 1 |
| PIxPI_138 | 1 | 1 | 1 |
| PIxPI_164 | 1 | 1 | 1 |
| PIxPI_165 | 1 | 1 | 1 |
| PIxPI_94 | 1 | H | 1 |
| PIxPI_134 | H | 1 | 1 |

| Genotypes | *usw329* (CNL) | *usw330* (NL) | *Pst* IT |
|---|---|---|---|
| PIxPI_2 | H | H | 1 |
| PIxPI_3 | H | H | 1 |
| PIxPI_7 | H | H | 1 |
| PIxPI_13 | H | H | 1 |
| PIxPI_14 | H | H | 1 |
| PIxPI_18 | H | H | 1 |
| PIxPI_19 | H | H | 1 |
| PIxPI_20 | H | H | 1 |
| PIxPI_22 | H | H | 1 |
| PIxPI_23 | H | H | 1 |
| PIxPI_26 | H | H | 1 |
| PIxPI_33 | H | H | 1 |
| PIxPI_35 | H | H | 1 |
| PIxPI_38 | H | H | 1 |
| PIxPI_39 | H | H | 1 |
| PIxPI_40 | H | H | 1 |
| PIxPI_41 | H | H | 1 |
| PIxPI_43 | H | H | 1 |
| PIxPI_44 | H | H | 1 |
| PIxPI_48 | H | H | 1 |
| PIxPI_51 | H | H | 1 |
| PIxPI_52 | H | H | 1 |
| PIxPI_53 | H | H | 1 |
| PIxPI_54 | H | H | 1 |
| PIxPI_56 | H | H | 1 |
| PIxPI_57 | H | H | 1 |
| PIxPI_58 | H | H | 1 |
| PIxPI_62 | H | H | 1 |
| PIxPI_63 | H | H | 1 |
| PIxPI_64 | H | H | 1 |
| PIxPI_65 | H | H | 1 |
| PIxPI_66 | H | H | 1 |
| PIxPI_70 | H | H | 1 |
| PIxPI_71 | H | H | 1 |
| PIxPI_78 | H | H | 1 |
| PIxPI_80 | H | H | 1 |
| PIxPI_82 | H | H | 1 |
| PIxPI_84 | H | H | 1 |
| PIxPI_90 | H | H | 1 |
| PIxPI_91 | H | H | 1 |

| Genotypes | *usw329* (CNL) | *usw330* (NL) | *Pst* IT |
|---|---|---|---|
| PIxPI_93 | H | H | 1 |
| PIxPI_98 | H | H | 1 |
| PIxPI_97 | H | H | 1 |
| PIxPI_99 | H | H | 1 |
| PIxPI_111 | H | H | 1 |
| PIxPI_113 | H | H | 1 |
| PIxPI_119 | H | H | 1 |
| PIxPI_126 | H | H | 1 |
| PIxPI_129 | H | H | 1 |
| PIxPI_132 | H | H | 1 |
| PIxPI_136 | H | H | 1 |
| PIxPI_143 | H | H | 1 |
| PIxPI_144 | H | H | 1 |
| PIxPI_148 | H | H | 1 |
| PIxPI_150 | H | H | 1 |
| PIxPI_156 | H | H | 1 |
| PIxPI_160 | H | H | 1 |
| PIxPI_161 | H | H | 1 |
| PIxPI_162 | H | H | 1 |
| PIxPI_171 | H | H | 1 |
| PIxPI_172 | H | H | 1 |
| PIxPI_29 | 0 | 0 | 2 |
| PIxPI_47 | 0 | 0 | 2 |
| PIxPI_69 | 0 | 0 | 2 |
| PIxPI_79 | 0 | 0 | 2 |
| PIxPI_103 | 0 | 0 | 2 |
| PIxPI_141 | 0 | 0 | 2 |
| PIxPI_173 | 0 | 0 | 2 |
| PIxPI_15 | 0 | H | 2 |
| PIxPI_30 | 0 | H | 2 |
| PIxPI_46 | 1 | 1 | 2 |
| PIxPI_89 | 1 | 1 | 2 |
| PIxPI_118 | 1 | 1 | 2 |
| PIxPI_127 | 1 | 1 | 2 |
| PIxPI_131 | 1 | 1 | 2 |
| PIxPI_168 | 1 | 1 | 2 |
| PIxPI_100 | H | 1 | 2 |
| PIxPI_12 | H | H | 2 |
| PIxPI_17 | H | H | 2 |
| PIxPI_24 | H | H | 2 |

| Genotypes | *usw329* (CNL) | *usw330* (NL) | *Pst* IT |
|---|---|---|---|
| PIxPI_28 | H | H | 2 |
| PIxPI_42 | H | H | 2 |
| PIxPI_45 | H | H | 2 |
| PIxPI_50 | H | H | 2 |
| PIxPI_55 | H | H | 2 |
| PIxPI_61 | H | H | 2 |
| PIxPI_67 | H | H | 2 |
| PIxPI_73 | H | H | 2 |
| PIxPI_76 | H | H | 2 |
| PIxPI_77 | H | H | 2 |
| PIxPI_83 | H | H | 2 |
| PIxPI_88 | H | H | 2 |
| PIxPI_96 | H | H | 2 |
| PIxPI_107 | H | H | 2 |
| PIxPI_109 | H | H | 2 |
| PIxPI_112 | H | H | 2 |
| PIxPI_115 | H | H | 2 |
| PIxPI_117 | H | H | 2 |
| PIxPI_122 | H | H | 2 |
| PIxPI_125 | H | H | 2 |
| PIxPI_128 | H | H | 2 |
| PIxPI_137 | H | H | 2 |
| PIxPI_147 | H | H | 2 |
| PIxPI_149 | H | H | 2 |
| PIxPI_151 | H | H | 2 |
| PIxPI_152 | H | H | 2 |
| PIxPI_153 | H | H | 2 |
| PIxPI_154 | H | H | 2 |
| PIxPI_163 | H | H | 2 |
| PIxPI_166 | H | H | 2 |
| PIxPI_25 | NA | H | 2 |
| PIxPI_31 | 0 | 0 | 3 |
| PIxPI_121 | 0 | 0 | 3 |
| PIxPI_133 | 0 | 0 | 3 |
| PIxPI_146 | H | H | 3 |
| PIxPI_106 | 0 | 0 | 4 |
| PIxPI_123 | 0 | 0 | 5 |
| PIxPI_145 | 0 | 0 | 5 |
| PIxPI_175 | H | H | 5 |
| PIxPI_158 | NA | H | 5 |

| **Genotypes** | ***usw329* (CNL)** | ***usw330* (NL)** | ***Pst* IT** |
|---|---|---|---|
| PIxPI_114 | 0 | 0 | 6 |
| PIxPI_104 | H | H | 6 |
| PIxPI_60 | 0 | 0 | 7 |
| PIxPI_116 | 0 | 0 | 7 |
| PIxPI_105 | H | 0 | 7 |
| PIxPI_34 | 0 | 0 | 8 |
| PIxPI_87 | 0 | 0 | 8 |
| PIxPI_120 | H | H | 8 |
| PIxPI_1 | 0 | 0 | 9 |
| PIxPI_11 | 0 | 0 | 9 |
| PIxPI_95 | 0 | 0 | 9 |
| PIxPI_140 | 0 | 0 | 9 |
| PIxPI_159 | 0 | 0 | 9 |
| PIxPI_176 | 0 | 0 | 9 |
| PIxPI_157 | NA | NA | 9 |

**Table S12:** Subset of the Global Durum Wheat Panel (GDP) screend using the *Yr84* (*CNL*) associated *usw314* KASP marker. None of the accessions carry functional PI 487260 *CNL* allele.

| **Code** | **Experimental Name** | ***Yr84* (PI 487260 *CNL*)** |
|---|---|---|
| GDPv2-019 | ESDCB-2015/2016-54 | - |
| GDPv2-031 | Calero | - |
| GDPv2-034 | BUCK_CANDISUR | - |
| GDPv2-038 | BUCK_C (NO_COMMERCIAL) | - |
| GDPv2-046 | BONAERENSE-INTA-CARILO | - |
| GDPv2-055 | CBW_08112 | - |
| GDPv2-062 | DAKTER | - |
| GDPv2-069 | MERIDIANO | - |
| GDPv2-113 | 1A.1D_5+10-6/3*MOJO//RCOL | - |
| GDPv2-124 | IDSN46-7014 | - |
| GDPv2-132 | IDSN46-7063 | - |
| GDPv2-137 | IDSN46-7093 | - |
| GDPv2-141 | IDSN46-7157 | - |
| GDPv2-154 | DAWRyT-0304 | - |
| GDPv2-160 | F11_00257=Ouassara | - |
| GDPv2-162 | IDON37-010 | - |
| GDPv2-177 | Icarukus2012 | - |
| GDPv2-217 | Extradur | - |
| GDPv2-230 | 1A.1D5+106/2*WB881//1A.1D5+106/3*MOJO/3/BISU1 /PATKA3 | - |
| GDPv2-235 | MESSAPIA | - |
| GDPv2-238 | Levante | - |
| GDPv2-244 | GALLARETA | - |
| GDPv2-263 | TENSIFT1 | - |
| GDPv2-270 | Lesina | - |
| GDPv2-283 | Biskri/Bouteille | - |
| GDPv2-286 | Borgia | - |
| GDPv2-288 | JM-3987 | - |
| GDPv2-296 | Arcobaleno | - |
| GDPv2-311 | Tetradur | - |
| GDPv2-325 | CORE | - |
| GDPv2-326 | Monastir | - |
| GDPv2-329 | Tirex | - |
| GDPv2-348 | Avonlea | - |
| GDPv2-359 | DYLAN | - |
| GDPv2-360 | Exeldur | - |
| GDPv2-368 | Neodur | - |
| GDPv2-380 | Bronte | - |
| GDPv2-389 | ELS_6404-127 | - |
| GDPv2-392 | Huguenot | - |
| GDPv2-400 | Apulicum | - |

| Code | Experimental Name | *Yr84* (PI 487260 *CNL*) |
|---|---|---|
| GDPv2-416 | Loukos2 | - |
| GDPv2-419 | Balloran | - |
| GDPv2-420 | Outrob5 | - |
| GDPv2-444 | Maci115 | - |
| GDPv2-448 | Margherita | - |
| GDPv2-456 | EL4X_27 | - |
| GDPv2-457 | EL4X_28 | - |
| GDPv2-465 | EL4X_69 | - |
| GDPv2-479 | EL4X_104 | - |
| GDPv2-482 | EL4X_122 | - |
| GDPv2-547 | EL4X_489 | - |
| GDPv2-550 | ENGB-Sohag-12 | - |
| GDPv2-553 | Ginchi | - |
| GDPv2-577 | Karabalykskaya_chernokolosaya_20 | - |
| GDPv2-583 | DW-SR-FIGS027 | - |
| GDPv2-584 | DW-SR-FIGS049 | - |
| GDPv2-590 | DW-SR-FIGS119 | - |
| GDPv2-617 | FIGSDWHOTCLD009 | - |
| GDPv2-618 | FIGSDWHOTCLD021 | - |
| GDPv2-637 | Susah | - |
| GDPv2-642 | SSD_006 | - |
| GDPv2-658 | SSD_067 | - |
| GDPv2-660 | SSD_092 | - |
| GDPv2-666 | SSD_115 | - |
| GDPv2-683 | SSD_244 | - |
| GDPv2-694 | SSD_294 | - |
| GDPv2-700 | SSD_326 | - |
| GDPv2-701 | SSD_336 | - |
| GDPv2-714 | SSD_470 | - |
| GDPv2-735 | SCORSONERA_(Saragolla) | - |
| GDPv2-736 | CANU_ARDESU | - |
| GDPv2-741 | MINGCHINOR | - |
| GDPv2-743 | MAKUZ-3 | - |
| GDPv2-748 | Kyperounda | - |
| GDPv2-754 | Akbasak | - |
| GDPv2-758 | Abyssinicum | - |
| GDPv2-759 | Abu_Fashit_NPGS | - |
| GDPv2-760 | T-2 | - |
| GDPv2-804 | PI341800 | - |
| GDPv2-809 | PI585018 | - |
| GDPv2-811 | CItr14629_TR02ID_SD | - |
| GDPv2-813 | BYAL258027 | - |
| GDPv2-816 | PI221423_Novo | - |
| GDPv2-817 | CItr14629_ELS 6404-75-3 | - |

| **Code** | **Experimental Name** | ***Yr84*** **(PI 487260 *CNL*)** |
|---|---|---|
| GDPv2-818 | PI115816 | **-** |
| GDPv2-819 | PI572849 | **-** |
| GDPv2-858 | PI470944 | **-** |
| GDPv2-859 | PI470945 | **-** |
| GDPv2-860 | PI352324 | **-** |
| GDPv2-883 | MG3430 | **-** |
| GDPv2-887 | MG4360 | **-** |
| GDPv2-888 | MG4366/2 | **-** |
| GDPv2-902 | MG5313 | **-** |
| GDPv2-957 | MG4411 | **-** |
| GDPv2-958 | MG4413 | **-** |
